# The hydrophobicity of peptides limits their availability to bind to interacting domains

**DOI:** 10.64898/2026.09.21.752983

**Authors:** Pascale Lemieux, Xavier Barbeau, Soham Dibyachintan, Alexandre K Dubé, Patrick Lagüe, Christian R Landry

## Abstract

A challenge in synthetic biology is to transfer the knowledge acquired *in vitro* to *in vivo* applications. For instance, binding affinity between peptide binding domains (PBDs) and their binding peptides is usually measured in simple buffers and in the absence of the cell machinery responsible for synthesis and proteostasis. Consequently, there is often a discrepancy between interaction quantified *in vitro* and *in vivo*. To identify strongly binding PBD-peptide pairs for *in vivo* applications, we aim to better understand the determinants of synthetic interaction in living yeast cells. We designed libraries of peptides expected to bind to PBDs based on data collected by phage-display. Using an *in vivo* protein interaction assay, we assessed the interaction strength of >6,500 PBD-peptide variants in yeast. We found that most substitutions disrupt binding, but that 3.4% increased binding strength to levels higher than expected from the phage-display data. We asked if, by using our mutational data, it was possible to optimize peptide sequences to gain even more binding strength and show additivity effect up to four substitutions. We found that PBD specificity is consistent between *in vitro* and *in vivo* assays. We hypothesized that *in vivo* peptide abundance could contribute to the discrepancy between the PCA and the phage-display results. We performed assays to detect *in vivo* peptide availability, reporting on peptide abundance, and we found that peptides with a stronger binding also show a higher availability in cells. We show that peptide availability, which seems mainly defined by peptide hydrophobicity, is key for the design of strong synthetic interactions.

## Introduction

Interactions between proteins are driving cell functions. Many protein-protein interactions (PPIs) are formed by the recognition of peptide motifs, present in intrinsically disordered regions, by peptide binding domains (PBDs). It is estimated that 10 000 PBDs are encoded in the human proteome, which have different specificities and could bind to more than 100 000 peptide motifs, contributing massively to the interactome (Tompa et al., 2014). The PPIs between PBD and peptide are transient, specific and involved in diverse cellular functions such as in signalling cascades, cytoskeleton organisation and condensate formations (Gilat et al., 2026; Holehouse & Kragelund, 2024). Over the last decades, massive efforts to characterize the binding preference of PDBs were done since many disease-causing mutations and virus infections are known to affect PPIs (Mészáros et al., 2021; Mihalič et al., 2023; Rrustemi et al., 2024).

These efforts led to the classification of PBDs in families that recognize conserved peptide motifs, such as SH3 domains and PDZ domains which bind respectively proline-rich and hydrophobe C-terminal motifs (Tonikian et al., 2008, 2009). Generally, the PBD-peptide motif interactions are studied by isolating the motif (< 10 amino acids) from its protein context and only characterize the PPI (Nardella et al., 2023; Tahti et al., 2023). The ‘simplicity’ of the PPI interface and the intrinsic binding preferences of PBDs makes PPIs a good model to study the effects of mutations on binding stability, which led to the development of exhaustive databases describing PPI affinities and binding preferences obtained *in vitro* (Kumar et al., 2024; Madhu et al., 2026; Teyra et al., 2020).

Using data mostly obtained *in vitro* on PBD binding to guide protein design, research groups were able to engineer orthogonal synthetic PPIs, such as synthetic scaffolds or binding competitive inhibitors, by expressing exogenous PBD and/or peptide in a cell host (Baek et al., 2013; Dueber et al., 2009; Guo et al., 2021). However, it is arduous to design these synthetic PPIs with consistency since the functions of exogenous proteins can be altered by the cell host (Cohen & Pielak, 2017). Indeed, exogenous proteins might be degraded or the proper folding and post-translational modifications might be inaccessible by the cell expression machinery (Zhang et al., 2007). Endogenous proteins could also interfere with the synthetic interactions by sequestering one or both of the interaction partners. The reciprocal effects could also occur whereby exogenous proteins would interfere with essential functions of the cells and lead to toxicity (Zarrinpar et al., 2003). Such factors might prevent an exogenous protein from performing its designed functions in cells. Thus, there is a need to better understand how predictive PPIs characterized in *vitro* data are for *in vivo* system design and what are the causes of the observed discrepancies.

Recently PBD functions and the residues involved in their binding preferences were thoroughly studied *in vivo* (Dibyachintan et al., 2025; Dionne et al., 2021; Faure et al., 2022), but their binding peptides received much less attention, with few exceptions (Jordan et al., 2025; Madhu et al., 2026). Peptide motifs generally do not have stable folds and can signal for localisation and degradation which can affect their abundance in cells (Tompa et al., 2014). A peptide which is degraded rapidly would not be of use in synthetic PPIs since the designed function would be limited by low abundance. Therefore, it is important to understand if the peptides bound strongly by PBDs *in vitro* have the same properties as the ones leading to strong PPIs in cells and how peptide abundance affects PPI strength.

Another important feature of synthetic PPIs is their orthogonality within a synthetic circuit. Orthogonality allows the specific recruitment of proteins and prevents cross talk in the designed circuit. In order to avoid non-desired PPIs, most synthetic circuits used PBDs from different families, assuming PBDs from the same family would have too similar binding preferences to allow orthogonality (Dueber et al., 2009). However, a growing body of literature suggests that the flanking residues of a motif contributes to binding affinity and specificity (Jordan et al., 2025; Kliche et al., 2024) and that PBDs from the same family can have different binding preferences (Tonikian et al., 2008, 2009). Thus, there is a need to understand how residues outside of the core motif influence the binding preferences of PBDs from the same family and whether they could allow orthogonal PPIs in cells.

In this study, we first compared the data characterizing the binding preferences of PBDs obtained in phage-display experiments with PPI measurements using peptide libraries in *Saccharomyces cerevisiae*. Second, we wanted to better understand if PBDs from the same family would maintain specificity by characterizing their PPI strength to libraries of each of their favored ligands in living cells. Third, we asked if we could design even stronger binders to PBDs in yeast using our mutational data. Then, we modelled PPIs *in silico* to assay the contribution of binding energies to PPI strength observed *in vivo*. Finally, we hypothesised that *in vivo* peptide abundance could explain discrepancies with *in vitro* expectations. Thus, we characterized the role of peptide availability, reporting on peptide abundance, on the PPI strength detected *in vivo*.

## Materials and Methods

Details on synthetic DNA, strains, plasmids, PCR mixes and cycles and materials are listed in Supplementary File 1 – Supplementary Tables.

### Culture media

The complex medium used for yeast transformation and growth was YPD [1 % yeast extract, 2 % tryptone, 2 % glucose, and 2 % agar (for solid medium)] to which we added nourseothricin (NAT, Cedarlane Labs) or hygromycin (HYG, Bioshop Canada) at final concentrations of 100 and 250 μg/ml, respectively. Synthetic medium [SC, 0.174 % yeast nitrogen base without amino acids and without ammonium sulfate, 2 % glucose, 0.1 % MSG, and 2 % agar (for solid medium)] depleted of uracil or leucine (-ura, –leu) were also used for yeast growth and selection. SC medium adjusted to pH 6 was buffered adding 0.6 % NaOH and 1 % succinic acid. All yeast cultures were incubated at 30 °C. Medium for bacterial growth was 2YT [1 % yeast extract, 1.6 % tryptone, 0.2 % glucose, 0.5 % NaCl and 2 % agar (for solid medium)] with ampicillin (AMP, Bioshop Canada) at final concentration of 100 μg/ml. All bacterial cultures were incubated at 37°C, under 250 r.p.m. agitation for liquid cultures. DHFR PCA selection condition was liquid synthetic medium (PCA medium, 0.67 % yeast nitrogen base without amino acids and without ammonium sulfate, 2 % glucose, 100mL drop-out without adenine and uracil) containing 200 μg/ml methotrexate (MTX, BioBasic) diluted in dimethyl sulfoxide (DMSO, Bioshop Canada). Drop out recipes are described in Table S8.

### Background strain construction

PL0002 strain was constructed based on the design of (Aranda-Díaz et al., 2017) by inserting the GEM transcription factor at the *leu2* locus of the BY4742 strain. The pHES839 plasmid was digested with PmeI and the resulting fragments were transformed using standard lithium acetate yeast transformation (Gietz & Schiestl, 2007) and plated on selective medium (SC-leu). PL0002, which now has a *LEU2*::GEM genotype, was validated by colony PCR (CLOP398-A3 to D3).

### Plasmid construction

The destination plasmid (pGD110), kindly donated by Guillaume Diss (Faure et al., 2022), expresses the murine dihydrofolate reductase fragments, DHFR F[1,2] and DHFR F[3], abbreviated as F[1,2] and F[3], on opposite DNA strands. Both fragments are under the control of the CYC promoter and CYC terminator and fused to a sequence that encodes a flexible linker in C-terminal (F[1,2]-(GGGGS)_4_, F[3]-(GGGGS)_3_-(GGGAS)_1_). The pGD110-variants encoding a PBD in C-terminal of F[1,2] were retrieved from a previous study (Lemieux et al., 2026) (Table S3). These pGD110 variants and pGD110 were digested with HindIII-HF (NEB) and purified using magnetic beads with a 1:1 ratio beads:dna (AMPure XP Beads, Beckman Coulter) for the insertion of sequences in frame with the F[3] in N-terminal. The peptide sequences were inserted downstream of F[3] paired with a F[1,2]-PBD or alone. The peptide sequences were ordered as oligonucleotides (IDT) and used to amplify pGD110 (PCR mix and cycle 1, Oligo F: C) and the purified amplicons were used for Gibson assembly. Gibson assembly was performed using 50 ng of digested plasmids with an insert molar ratio of 1:3 (backbone: insert) in a 10 μL reaction volume (Gibson et al., 2009). The reactions were incubated at 50 °C for 60 minutes and 5 μL of the reactions were transformed in MC1061 chemo-competent cells. All transformations were plated on solid 2YT+AMP. The same cloning strategy was used to insert peptide sequences in pPL9 which encodes the mEGFP (abr. GFP)-(GGGGS)_3_-(GGGAS)_1_ under the control of the CYC promoter and terminator (Lemieux et al., 2026). All the synthetic DNA used for the plasmid construction is described in Tables S1 and S2. All plasmids that are listed in Table S3 were validated by colony PCR and most, by Sanger sequencing (CHUL sequencing platform).

### Strain constructions – Landing Pad with F[1,2]-PBD and F[3]-peptide

The pGD110-variants encoding the PBD 363, PBD 366 and PBD 385 were used as DNA template for the PCR amplification of the PBD sequence. They were amplified using the standard PCR reaction and cycle with primers adding 40-pb homology arms to the *GAL1* locus in 5’ and to the NATMX marker in 3’ of the F[1,2]-linker-PBD-CYCt (CLOP197-G6/CLOP389-H12). The NATMX marker was amplified from pAG25-DHFR F[1,2] (Tarassov et al., 2008) using the standard PCR mix and cycle (CLOP389-G12/CLOP312-G8) adding 40-pb homology to the CYCt in 5’ and to the *GAL1* locus in 3’. Then, a fusion PCR was performed for which a 1:50 dilution of the PCR fragments of the first amplifications served as the DNA template (PCR mix and cycle 2, CLOP197-G6/CLOP312-G8). The resulting F[1,2]-PBD-CYCt-NATMX fragments were transformed by homologous recombination in PL0002 using standard lithium acetate yeast transformation (Gietz & Schiestl, 2007) and plated on selective medium (YPD+NAT), resulting in the strains PL0122 to PL0125 (Table S4). The correct genomic integrations were validated by colony PCR (CLOP248-B7/CLO1-61) and Sanger sequencing.

The same strategy was used to insert the F[3]-peptide-CYCt-HPHMX at the *GAL1* locus (PL0077 to PL0121, Table S4). The pGD110-variants encoding the peptide sequences of interest and pGD110 were used as DNA template and the F[3]-peptide-CYCt fragments were amplified (CLOP197-B7/CLOP370-E7). The pAG32-DHFR F[3](Tarassov et al., 2008) was used as a DNA template for the amplification of the HPHMX marker (CLOP312-G8/CLOP370-F7). Then, a fusion PCR was performed resulting in the F[3]-peptide-CYCt-HPHMX fragments with 40-pb homology arms to the *GAL1* locus (PCR mix and cycle 2, CLOP312-G8/CLOP197-B7). The yeast transformations were done in the PL0001 background using YPD+HYG as a selective medium and validations were performed as described above (CLOP248-B7/CLO1-57).

### Yeast mating and diploid selection

Haploid yeast strains encoding either the NAT or HYG resistance cassettes were grown on solid selective media and resuspended in YPD. *MAT**a*** and *MATα* strains were mixed by inoculating 25 μL of each resuspension in 200 μL YPD and incubated overnight at 30 °C. Then, the cultures were spotted or striked on solid medium for diploid selection (YPD+NAT+HYG) and incubated at 30 °C for two days.

### Small-scale DHFR PCA assay and analysis

The DHFR PCA protocol is based on Tarassov et al. 2008. We transformed the BY4741 strain with pGD110-variants expressing PBD and peptide sequences, single or paired, and we used DHFR PCA to quantify the PPIs (Gietz & Schiestl, 2007). We also performed a DHFR PCA to quantify the binding availability (Levy et al., 2014) of each PBD and peptide using a previously published strain (PL0017). This strain expresses from its genome a yellow fluorescent protein fused to the F[1,2] from the *HO* locus (Lemieux et al., 2026). We transformed PL0017 strains with pGD110-variants encoding a peptide without a PBD. Yeast strains transformed with pGD110-variants were grown in liquid synthetic medium (SC-ura), then diluted to an optical density (OD) at 595 nm of 0.1 in DHFR PCA selection medium and incubated in transparent polystyrene 96-well plates (Greiner Bio-One). Growth rates from three independent replicates were taken without shaking by measuring the OD at 595 nm in 15-minute intervals over two days of incubation at 30 °C in a BioTek Epoch 2 plate reader (Agilent). The median of the first five measurements was subtracted from all the following measurements to correct for variation in initial cell inoculate density. Empirical area under the curve (corrected AUC) was computed using the first 40 hours of growth. The corrected AUC metric was chosen over the derivative growth rate because the corrected AUC showed weaker background signal.

The same protocol was used for the diploid strains expressing from the *GAL1* locus a F[3]-peptide and a F[1,2]-PBD, with the exception of the media used. Diploid strains were grown in liquid synthetic medium overnight (SC+NAT+HYG pH 6) and the growth curves were performed in PCA medium containing MTX and 8 nM β-estradiol (Sigma-Aldrich).

### Double mutant peptide libraries design and construction

From a previous study that used small-scale DHFR PCA PPI quantification between a PBD and peptides (Lemieux et al., 2026), we selected the PBD-peptide pairs which showed PCA signal (corrected AUC >25) validating that the PPI occurs *in vivo* (three PDZ and four SH3 domains). For each PBDs, we used their previously characterized position weight matrices (Teyra et al., 2020) to extract a PWM_max_ peptide which encodes the most favored amino acid for binding at each position as defined by phage-display (Figure S1). Then, we designed a library with two degenerated positions using the PWM_max_ peptide as background sequence which will be tested against their respective PBD. The positions to degenerate were selected based on the PWM where each amino acid at each position has a score from 0 to 20. A score of 20 indicates an amino acid essential for binding. For each PWM, we kept the amino acids with a score higher than 15 because we expected these residues to be essential for the peptide recognition at those positions. The two positions encoding an amino acid with the next highest scores below 15 were degenerated with a NNK codon. The remaining positions of the peptide were kept as the top scoring residues, creating 400 double peptide mutants of the PWM_max_ peptide for each PBD encoded in 1024 DNA sequences. We codon-optimized the remaining positions of peptide sequences for expression in *S. cerevisiae* and synthesized them as oligonucleotides (Table S2, IDT).

We used the same cloning strategy for the library construction as previously described for the pGD110-variants construction, but with enough replication to obtain a 10X coverage over the sequence diversity. We performed the degenerated peptide amplification in duplicate with PCR mix and cycle 1 (Oligo F: <u>D</u>). The amplicons were purified using magnetic beads (1:1 beads:DNA ratio, AMPure XP Beads, Beckman Coulter). The degenerated sequences were inserted in pGD110 paired with their respective PBD or alone. The Gibson assembly was executed in four replicates for each degenerated peptide. Each reaction was transformed in MC1061 chemo-competent cells twice and plated on selective medium (2YT+AMP). We used 5 mL of 2YT liquid medium to collect the clones by scraping the transformation Petri dishes and we measured the OD for each scraping. The eight scrapings of one degenerated peptide were pooled proportionally to the scraping with the highest OD, avoiding the dilution of sequence diversity i.e. adding less volume from the less efficient clonings/transformation to the pool. We used 750 μL of each pool to extract the plasmid libraries (Presto Mini Plasmid Kit, Geneaid Biotech Ltd). We modified the kit protocol to increase plasmid extraction efficiency with samples of high cell density. We used 300 μL of PD2 buffer for cell lysis and 450 μL of PD3 buffer for neutralization.

### DHFR PCA bulk competition

The DHFR bulk competition protocol is based on Dubé et al. 2022 (Dubé et al., 2022). The plasmid libraries encoding for pairs of PBD-degenerated peptides were transformed in BY4741 for the PPI assay and plasmid libraries encoding for degenerated peptides alone were transformed in PL0017 (Lemieux et al., 2026) for the binding availability assay. We transformed each plasmid library to obtain a 10X coverage over the sequence diversity and we plated all transformations on selective medium (SC-ura) (Gietz & Schiestl, 2007). We used 5 mL of SC-URA to collect the clones by scraping the Petri dishes and we measured the OD of each scraping. The scrapings of the same libraries were pooled proportionally to the scraping with the highest OD and conserved in glycerol stock. Reference pGD110-variants (Table S3) were freshly transformed in BY4741 or PL0017 and plated on selective medium (SC-ura). The reference sequences chosen were selected based on the previous characterization by growth curve analysis in the PPI and availability assay and cover the sensitivity range of the assays including the maximal (ref_max_) and minimal (ref_min_) signals obtained previously (Table S3) (Lemieux et al., 2026).

We inoculated cultures in 3 mL of selective medium (SC-ura) with 50 μL of the library glycerol stocks in triplicate and grew them overnight in a deep-well plate at 30 °C under 400 rpm shaking. We also inoculated cultures in triplicates using the fresh transformation of the reference pGD110-variants (Table S3). We measured the OD of each culture. We pooled in triplicate each library with equal OD to obtain a similar frequency of each peptide library in the pool. We pooled in triplicates the reference pGD110-variants, from independent cultures, using equal OD units from each culture. The pGD110-variants reference strains were spiked in the pooled library at 5 % frequency. We collected 5 OD units of each pooled replicate as an initial time point (T0) and the cell pellets were stored at –80 °C. Each replicate of the pool is diluted at 0.15 OD in 15 mL (Falcon 50 mL tube) of PCA selection medium and incubated at 30 °C under 250 rpm shaking. Each replicate was grown to 2.5 OD (∼4-5 generations) where 5 OD units were collected (T1) and cell pellets stored at –80 °C. We diluted each replicate at 0.15 OD in fresh PCA selection medium and the same procedure was followed when the final time point was reached (2.5 OD, ∼4-5 generations, T2).

### Double mutant libraries bulk competition sequencing and analysis

The cell pellets (T0, T1 and T2) collected during the bulk competition were used to extract the plasmid libraries with the Zymoprep Yeast Plasmid Miniprep Kits (Zymo Research). The plasmid extraction protocol was modified by increasing the amount of zymolyase to 30 U per sample. We also added a centrifugation step (5 minutes at 15 000 rpm) to improve the separation of the cell debris from the supernatant. The eluted plasmids were used as template DNA for the first PCR (PCR1) for library preparation (PCR mix and cycle 3). The PCR1s were performed in duplicates using one oligo pair for all samples of the same time point.

The duplicated reactions were pooled and purified using magnetic beads (AMPure XP Beads, 1:1 ratio beads:DNA). The purified PCR1 products were used as DNA template for the second PCR (PCR2) adding the Illumina adaptors and Nextera index. The PCR2s were also performed in duplicates following the PCR mix and cycle 4 (Oligo F: <u>G</u>, Oligo R: <u>H</u>, Table S6).

The duplicated PCR2s were pooled and purified using magnetic beads (AMPure XP Beads, ratio DNA:beads 1:1). All samples were pooled together with 50 ng each and sent to sequencing (NovaSeq paired-end 150 pb, CHUL sequencing platform). We obtained >23 millions reads for all samples which represents an overall coverage of >175X.

We used FastQC v0.11.9 to visualize the reads’ quality for each sample. We trimmed the reads with Cutadapt v3.5 in paired-end mode using the constant homology regions upstream and downstream of the sequences of interest ( -a ACTGTCACTGACGCGGGAGGTGGAGCTAGC…aagcttattagttatgtcacgcttacattcacgccctc –A gagggcgtgaatgt aagcgtgacataactaataagctt…GCTAGCTCCACCTCCCGCGTCAGTGACAGT). Pandaseq v2.11 was used to merge the reads with the minimal and maximal length fixed to 10 and 100 base pairs respectively (`-l 10 –L 100 –T 6`). Finally, vsearch v2.30.0 was used to count the occurrences of unique sequences. We assigned the reads to each peptide sequence by perfect match. We retrieved 20 million reads using this workflow.

### Second DHFR PCA bulk competition with single mutant and cross-specificity libraries

Following the results for the double mutants DHFR PCA bulk competition experiment which showed mutation additivity (Figure 2), we decided to test all single mutants of the PWM_max_ for three PBDs (363, 366, 385). We degenerated peptide positions, which were not already included in the double mutant library, with a NNK codon. The DNA sequences were codon optimized for expression in *S. cerevisiae*, and synthesized as oligonucleotides (Table S2, IDT). The peptide sequences were amplified (PCR mix and cycle 1, Oligo F: I) and inserted in pGD110 paired with their respective PBD or alone using the same method as for the double mutant libraries. We also wanted to test for binding specificity for the three selected PBDs, thus we inserted the double mutants and the single mutants of each peptide paired with the two other PBDs (ex. 363 peptide library is also inserted paired with PBD 366 and PBD 385). We pooled together 1.5 μg of each the single mutant library to create one library per pair of PBD-peptide libraries. We transformed the new plasmid libraries, single mutants and new double mutants, into the strains PL0017, when alone, and BY4741, when paired with a PBD, with a 10X coverage over the sequence diversity i.e. collecting ten times more colonies than individual sequence in the libraries. The yeast libraries were recovered and pooled as described for the double mutant libraries.

The cultures from the library glycerol stock were inoculated in triplicate as described above. We used the same reference pGD110-variants for the PPI and availability assays as described in the first bulk competition. For the availability assay of the single mutants, we pooled with equal OD values the single mutants libraries. For the cross-specificity competition, we pooled the single and double mutant cultures together relative to their sequence diversity, creating one pool per pair of PBD-peptide libraries. For the fitness assays in both strain background, we pooled the previous four PBD-peptide libraries (152, 246, 250, 299, in BY4741) with equal OD values, and also pooled all the double mutants libraries unpaired with a PBD (PL0017) with equal OD values. The same reference sequences as used in the previous bulk competition are spiked in the pools at a 5 % frequency. We also performed a fitness assay for the single mutant availability pool and the cross-specificity pools using MTX depleted PCA medium (DMSO). The bulk competitions are performed in the same way as described above. The different assays and the sequences included in each one are summarized in Table S6.

The plasmid extraction and PCR1s (PCR mix and cycle 3) were performed as described above. The PCR2s (PCR mix and cycle 4) to add the Nextera indexes and Illumina adapters were also done as described above using the oligonucleotides listed in Table S6 (Oligo F: <u>J,</u> Oligo R: <u>K</u>). The 72 samples were pooled relative to their sequence diversity with a minimum of 3.9 ng per sample. The DNA pool was sent to deep sequencing (NovaSeq paired-end 150 pb, CHUL sequencing platform) and we obtained 40 millions reads which was below the requested coverage (100X). Thus, we requested the pool to be re-sequenced with more coverage on the same machine and the combined number of reads from the two runs was 214 millions. It represents an average coverage of 1225X per sample (minimum coverage 66X, maximum coverage 2380X, Figure S6A). The reads processing followed the same pipeline as described above and 203 millions reads passed all the filters.

### PCA score analysis

To assign scores reporting the strength of the PPI or availability signal to each peptide sequence, we computed the log2 fold change in read frequency of DNA variants for each combination of pools, replicates and conditions. First, we added 1 to the read count of each sequence at the three timepoints in all samples. We kept individual DNA sequences when we counted >20 reads in T0 samples for at least two out of three replicates. If the number of reads was below 20 in one replicate at T0, the DNA sequence was removed from the replicate. Then, we computed the frequency of each peptide DNA sequence in their respective sample at each timepoint. We calculated the difference between T2 and T0 of log_2_(freqT) and normalised it with the number of generations between the timepoints resulting in a selection coefficient (s). We transformed the s by standardization relative to the reference sequences so that we could compare each pool together (Figure S2). The standardization resulted in normalised s value ranging from 0 to 1 which is the signal considered in the sensitivity range of the growth curve assays.

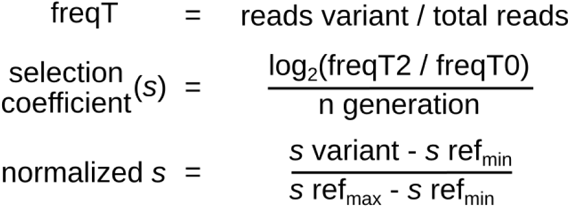

For the second DHFR PCA bulk competition, we filtered the normalised s value of each replicate to remove outliers. To do so, a generalized linear model was fitted between the comparisons of each replicate, i.e. R1 vs R2 & R1 vs R3 & R2 vs R3. The coefficient and the intercept of the models associated with their standard error were retrieved. The difference between the models maximal and minimal values of each parameter (ex. model 1 coef.+standard error – model 3 coef.-standard error) were summed to obtain a tolerance threshold of variability between replicates. If the empirically determined threshold value was higher than 0.4, it was capped at 0.4 of normalised s difference. The difference in normalised s value between the replicate of one peptide DNA sequence was computed across the three replicates and, if the normalised s of one replicate was more different than the threshold value with the other two replicates, the problematic replicate was removed (replication index <2, Figure S6C, Figure S7).

Using the filtered normalised s values, we performed statistical tests, two-ways Welch t-test with a Benjamini-Hochberg correction (FDR), within each peptide family and each pool to compare every peptide sequences (which included the three technical replicates and could include different codons as internal replicates) to the PWM_max_ peptide. When the corrected p-values < 0.05, the peptide was identified as different from the PWM_max_ peptide and, whether it had a stronger or weaker PCA signal was determined by the sign of the Welch t-statistic (stronger > 0, weaker < 0). Finally, the PCA scores, PPI and availability, were obtained by computing the median of normalised s values across the replicates for each peptide variant (Figure S8, S10-S12). We compared the PCA scores of the sequences included in both bulk competitions and we obtained a good degree of replication (Figure 2C).

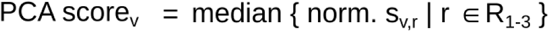

### Sequence selection and validation of bulk competition assays

In order to validate the large scale experiments testing peptide availability, binding strength and specificity, we selected 49 peptides present in the bulk competitions divided in three validation objectives. The first objective was to confirm the peptide variants with an increased PPI score vs the PWM_max_ peptides. Thus, we selected the top five peptide variants with a significantly stronger binding to their designed PBD (ex. peptide family 366 against PBD 366) in the second bulk competition and we also selected all other peptides with a PPI score > 0.9 even if they were not found to bind differently from the PWM_max_ peptide. The second objective was to validate the peptide variants which increase PPI score to their non-designed PBD (ex. peptide from family 366 with increased binding to PBD 385). All peptide variants with a significantly stronger binding vs PWM_max_ to non-designed PBDs were retrieved and filtered by PPI score > 0.46. The last objective was to validate the effect of stop codons in the PWM_max_ sequence on availability score. Thus, we selected the peptide variants with stop codons at position 2, 5, 8 and 10 in the 363, 366 and 385 PWM_max_ peptide sequences.

In addition to the peptide variants present in the bulk competition assay, we designed peptide sequences to further test the mutation additivity in PPI. For the PBD 363, PBD 366 and PBD 385, we selected the two double mutant sequences in their designed peptide family which had the top-2 significantly stronger PPI score. Then, we selected from the single mutant libraries two positions where the top scoring amino acid was different from the PWM_max_ peptide. From these positions, we chose the top-2 scored amino acids. We designed peptide variants with the top 2 amino acids at these positions in both double mutation backgrounds resulting in eight peptide variants per PBD (Figure 2I, Figures S12-14).

All the validation peptide variants were ordered as oligonucleotides (IDT, Table S2) and cloned in pGD110, paired with a PBD or alone, and/or in pPL9 as described above. Once validated by Sanger sequencing, the pGD110-variants encoding the validation sequences were transformed in either PL0017 or BY4741 for DHFR PCA small-scale measurements. The growth curves and analysis were done as described above.

### Structural modelling of PDZ-peptide complexes

PDZ-peptide complexes for PBD 363, PBD 366 and PBD 385 were predicted using AlphaFold3 (Abramson et al., 2024). Five models were generated for each complex. For each PBD, the five predicted structures were superposed on receptor Cα atoms and per-residue root mean square fluctuations (RMSF) were calculated across the resulting ensemble. Terminal regions comprising at least three consecutive residues with RMSF values > 1.5 Å were considered poorly converged and removed from the receptor sequence. The trimmed receptor sequences were resubmitted to AlphaFold3 and the top-ranked prediction (model 0) was retained for all subsequent analyses.

To assess model accuracy, the AlphaFold3 structures were superposed in PyMOL (Schrödinger, LLC) onto experimentally determined structures of the corresponding PDZ domains: PBD 363 (ZO-1 PDZ3, PDB 4Q2Q, bound to a phage-derived peptide) (Ernst et al., 2014), PBD 366 (Tiam1 PDZ, PDB 3KZD, apo; (Shepherd et al., 2010)) and PBD 385 (PDLIM4 PDZ, PDB 4Q2O, bound to a peptide ligand) (Ernst et al., 2014). Backbone Cα RMSD values calculated over aligned receptor residues were 0.81, 0.71 and 0.50 Å, respectively. Peptide conformations were evaluated for PBD 363 and PBD 385, whose crystallized ligands shared six and five aligned ordered residues with the modelled peptides, respectively. After superposition on the receptor alone, Cα RMSD values were 0.17 and 0.12 Å, indicating near-identical peptide conformations. For PBD 366, the crystallized ligand was insufficiently similar in sequence to permit an equivalent comparison.

### ΔΔG calculations

Peptide sequences and interaction scores were obtained from the datasets generated in this study. Changes in binding energy (ΔΔG) were estimated using FoldX version 5.1 (Delgado et al., 2025) with the parameter –-vdwDesign=2, using model 0 of each PBD-peptide complex as the reference structure. Prior to mutational analyses, reference structures were subjected to five successive rounds of RepairPDB. Total energy values reached a plateau after the fourth round and changed minimally thereafter.

Peptide variants were generated with BuildModel (--numberOfRuns=3), with all substitutions within a peptide introduced simultaneously. Interaction energies of the wild-type and mutant complexes were then evaluated using AnalyseComplex. ΔΔG values were calculated as the mutant minus wild-type interaction energy and averaged across the three independent FoldX runs. Positive ΔΔG values indicate mutations predicted to weaken binding, whereas negative values indicate stabilizing mutations.

Binding ΔΔG values were additionally estimated with the flex ddG protocol (Barlow et al., 2018), implemented through PyRosetta (Chaudhury et al., 2010) with the published ddG-backrub.xml script and the talaris2014 energy function. Wild-type and mutant states were constructed on backrub-sampled backbone ensembles using five ensemble members and 3500 backrub trials per variant. ΔΔG values were averaged across ensemble members and follow the same sign convention as above. Values are reported in Rosetta energy units (REU).

### Abundance measurements of GFP-peptide variants by flow cytometry

To measure the effect of peptides on the abundance of their fused protein, we cloned peptide sequences in the C-terminal of the GFP in the pPL9 backbone (Lemieux et al., 2026) as described above. The pPL9-variants were transformed in the strain BY4741 and plated on selective medium (SC-ura). The transformed strains were grown overnight in triplicates in SC-URA. Cultures were diluted in fresh media (OD ∼0.15) and grown to exponential phase (OD = 0.5-0.7), then they were washed and diluted in PBS to 0.05 OD for cytometry measurements. The fluorescent intensity of 5000 cells per replicates was measured using a Guava EasyCyte HT cytometer (excitation: blue laser, λ = 488 nm; detection: green channel, λ = 512 – 525 nm). The median fluorescent intensity of each replicate was computed and used to compare the pPL9-variants.

## Results

### DHFR PCA allows detection of stronger binders than the *in vitro* reference peptide

We retrieved from a previous study three PDZ and four SH3 domains that were shown to interact with peptides *in vitro* and in living yeast cells (Lemieux et al., 2026; Teyra et al., 2020). The PBD binding profiles were characterized previously by phage-display used to derive position weight matrices (PWMs) from random peptide libraries enrichment and to describe PBD binding preference (Teyra et al., 2020). A PWM attributes a score to each amino acid at each position of a peptide whose sum describes the binding preferences of a PBD (Figure S1). We used the PWMs to extract the favored amino acid at each position in the peptide to obtain the maximally binding one to each PBD *in vitro*, which we refer to as PWM_max_ peptides (Figure 1A).

**Figure 1.**
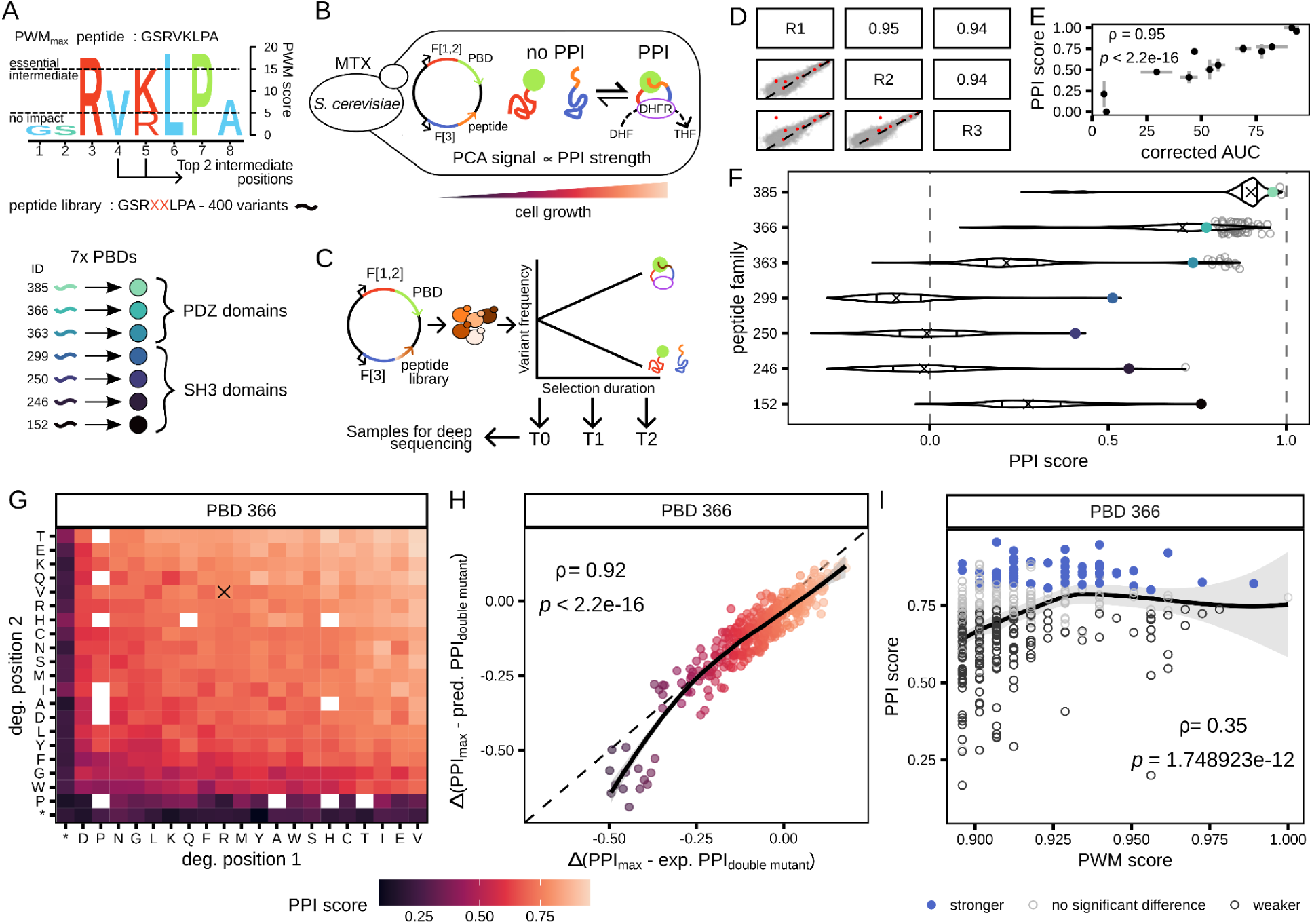
PPI strength characterization of the double mutant peptide library. **A.** The peptide library design was based on the PRM-DB PWMs where every amino acid at each position has an enrichment score from 0 to 20, where a score of 20 represents a position at which only one residue allows binding (Teyra et al., 2020). Here, an amino acid with a score above 15 is considered essential and a position where all amino acids have a score below 5 is considered to have minimal impact on binding. To enrich the peptide library with variants having an effect on binding, we selected the two positions with intermediate amino acid scores (5 < score < 15) as positions to degenerate from the PWM_max_ peptide. One library of double mutants was designed for each of the seven PBDs. **B.** Illustration of the DHFR PCA approach (Tarassov et al., 2008) to quantify binding strength between a PBD and a peptide *in vivo*. The PCA signal is to cell growth in selective medium, which depends on the amount of assembled DHFR that, itself, is proportional to the PPI strength between the F[1,2]-PBD and the F[3]-peptide (Freschi et al., 2013). **C.** Schematic representation of the DHFR PCA bulk competition framework (Diss & Lehner, 2018; Faure et al., 2022). The peptide library is encoded on plasmids, which are extracted before and after selection and deep sequenced. The variant frequency changes allow the quantification of competitive growth rates and thus, of relative PPI strength. **D**. Scatter plots representing the correlation between the normalised *s* from each replicate. The replicates are identified on the diagonal and the Spearman’s correlation coefficients are shown in the mirror quadrant to each scatter plot (p-values < 2.2e-16 for all comparisons). The red data points correspond to the reference sequences added in the libraries. **E.** Scatter plot showing the comparison between PPI scores obtained in competition and the PCA signal obtained from individual growth curve analysis (corrected AUC) for validation PBD-peptide pairs. Error bars represent the standard deviation obtained for each experiment performed in triplicate. The Spearman’s correlation coefficient and its p-value are shown. **F.** Violin plots representing the distributions of PPI scores for each peptide family. The colored data points are the PWM_max_ peptides scores and the empty data points represent the peptide variants with a significantly stronger PPI score than the PWM_max_ peptide. The median of each distribution is shown as a cross and the 25th and 75th quartiles are drawn as lines in the violins. The dashed lines at 0 and 1 represent the sensitivity range of the assay covered by the reference sequences. **G.** Heatmap of the PPI scores for each peptide variant in the 366 peptide family. The color scale represents the PPI score and the pairs of mutated residues are labelled on the axes. Amino acid order follows their average PPI score per position. The cross highlights the PWM_max_ peptide. White tiles are missing data. **H.** Scatter plot showing the correlation between the expected change in PPI scores of double mutants estimated from single mutant data (pred. PPI_double mutant_, y-axis) and the experimental changes observed for double mutants (exp. PPI_double mutant_, x-axis) in the 366 peptide family. The color scale represents the PPI score. The Spearman’s correlation coefficient and its p-value are labelled. A local regression is fitted on the data as a solid line. (exp. PPI_double mutant_, x-axis) and the difference in PPI score of the PWM_max_ peptide and the sum of PPI scores of single mutants (pred. PPI_double mutant_, y-axis)**I.** Scatter plot showing the relationship between the PPI and PWM scores of each peptide variant. The color indicates the peptide variants with significant differences when compared to the PWM_max_ peptide in the PCA assay. The Spearman’s correlation coefficient and its p-value are labelled. A local regression is fitted on the data as a solid line.

To design peptide libraries with a wide range of potential interaction strengths to the PBDs, we used the PWMs data to guide our design. We classified the positions of the PWMs according to their amino acids scores. We reasoned that impactful effects on binding could take place at positions with intermediate scores (15 >= aa score > 5) as the positions already favored (20 >= aa score > 15) would provide little gain and those where the amino acid identity does not influence binding (5 ≥ aa score) would have limited effects (Figure 1A). Using the PWM_max_ peptides as starting backgrounds, we degenerated the codons of 2 positions which encode amino acids with intermediate contributions to binding based on PWMs (Figure S1). We designed 7 double mutant libraries with 400 peptide variants each to test for changes in PPI strength. Each group of 400 peptide variants form a mutant peptide family named after their target PBD.

We used the dihydrofolate reductase protein fragment-complementation assay (DHFR PCA, Figure 1B) to characterize PPI strength between the PBDs and their peptide families (Tarassov et al., 2008). In this assay, the murine DHFR is split in two fragments, F[1,2] and F[3], which are fused to pairs of proteins and co-expressed in *S. cerevisiae*. In selective media conditions, the endogenous yeast DHFR is inhibited by methotrexate (MTX). When the DHFR fragment fused proteins interact, the two fragments restore the activity of the mDHFR which is MTX insensitive. Since the DHFR catalyses an essential reaction for yeast survival, its growth is relative to the amount of assembled mDHFR. Thus, the stronger the interaction between proteins fused to the DHFR fragments is, the faster the yeasts grow (Freschi et al., 2013). This assay has been widely used for the quantification of protein-protein (Dionne et al., 2021; Tarassov et al., 2008) and protein-peptide interaction including to measure the impact of mutations on binding (Figure 1B) (Bendel et al., 2024; Dibyachintan et al., 2025; Dionne et al., 2021; Diss & Lehner, 2018; Faure et al., 2022; Jordan et al., 2025; Lemieux et al., 2024).

We ordered each peptide family as synthetic DNA and inserted them in the plasmid in frame with F[3]-(GGGGS)_3_-(GGGAS)_1_ and paired with a F[1,2]-(GGGGS)_4_-PBD (Figure 1C). We performed a bulk competition experiment in which the peptide sequence acts as a DNA barcode to follow the frequency of each variant in the competition assay. The changes in sequence frequency in the competition depends on the amount of active mDHFR enabled by the PPI. We performed the bulk competition in triplicate and we collected cell pellets before and after the DHFR PCA selection. We extracted the plasmid library from these pellets and deep-sequenced them.

Using the DNA sequence frequencies, we computed PPI scores reporting the interaction strength between each peptide variant and its paired PBD. We included reference sequences in each library which showed a wide range of signals in small-scale DHFR PCA measurements (Lemieux et al., 2026) (Figure 1D). We used these references to normalize the change in read frequency between replicates (normalised *s*) (Figure S2B). The normalised *s* are highly correlated between replicates (ρ >= 0.94, p-value < 2.2e-16, Figure 1D). We performed statistical tests on the normalised *s*, combining the DNA sequences encoding the same peptide variant in the 3 replicates, in order to define which peptide variants show stronger or weaker binding than the PWM_max_ peptide in each peptide family (Wilcoxon’s tests, Benjamini-Hochberg correction, p-value < 0.05, Figure S2D & F). Then, we computed the median of normalised *s* from the 3 replicates of DNA sequence encoding the same peptide variant to obtain their PPI scores (Figure S2F, Figure S3). We compared the PPI scores and the growth curves measurements obtained in a previous study (Lemieux et al., 2026) and observed strong agreement between the two independent experiments, indicating highly reproducible results (Figure 1E, Spearman’s correlation, ρ = 0.95, p-value < 2.2e-16).

We observed that each peptide family has a similar distribution of effect on binding with their respective PBD (Figure 1F). The PWM_max_ peptides have higher PPI scores than the 75th quantile of the distribution for each family validating that the phage-display captured one of the strongest binders to a PBD. Most peptide variants disrupt the PPI as expected. However, we detected peptide variants with a significantly stronger interaction than the PWM_max_ peptide for 4 out of 7 PBDs (Figure 1F). From these 4 PBDs, 3 are PDZ domains (PBD 363, 366, 385), and they all show stronger PPI signal than the exogenous SH3 domains (PBD 246, 250, 299) tested (PBD 152 being the endogenous SH3 domain of Sho1p). The strong PPIs enabled by the PDZ domains make them good PBD candidates for further characterization and use in synthetic circuits in yeast.

When looking at each peptide variant, we observe that substitutions to amino acids with similar properties do not systematically lead to the same effect on PPI strength (Figure 1G, Figure S3). For example, a substitution in the PWM_max_ peptide 366 to an aspartic acid (D, negatively charged), at either position, leads to a decrease in PPI strength, while substitutions to glutamic acid (E) lead to an increase. The double mutants, DD and EE, have even more divergent effects, suggesting additivity of mutation effects on PPI strength. We therefore extended this analysis to all pairs of substitutions to see how common was additivity in PPI strength. Simple additivity would allow predicting binding strength of double mutants by summing that of the single mutants. Thus, we used the single mutant PPI scores in each peptide family to predict the double mutant impact on PPI strength. For all peptide families, we found highly significant correlations between our prediction and the PPI score of double mutants (Spearman’s correlation, ρ = 0.2 to 0.93, p-values < 1e-4, Figure 1F, Figure S4). The correlation coefficients are higher when the median PPI score of the peptide families is within the detection range of the assay (Spearman’s correlation, ρ = 0.6 to 0.93, p-values < 2.2e-16, Figure 1H, Figure S4). The inconsistent prediction accuracy could stem from single mutants already reaching the lower detection limit (PPI score < 0). For instance, combining two highly deleterious substitutions for the PPI would lead to twice the decrease in signal, assuming perfect additivity, which is not detectable in our experimental setup. For peptide variants leading to stronger PPIs, additivity of substitution effects is a good predictor of the PPI score since the upper limit of our assay was not reached by any peptide family (Figure 1F).

We used the PWMs data to predict the PPI scores obtained experimentally for each peptide variant. We calculated a PWM score by adding up the scores of each aa encoding the peptide variant at each position of the PWM and transformed it into a relative value compared to the PWM_max_ score. Since we based our libraries on PWM_max_ peptides (PWM score = 1), we expected a decrease in PWM score for each peptide variant, but we asked if this decrease would be more important for peptide variants leading to a decrease in PPI score. We found that PWM scores are not always correlated with PPI scores (Spearman’s correlation, ρ = –0.083 to 0.6, p-values = 0.77 to 2.2e-16, Figure S5), with peptides showing stronger PPI spread throughout the distribution of PWM scores (Figure 1I). The discrepancy between *in vivo* and *in vitro* scores reveals that *in vitro* experiments can miss the optimal peptide motifs for the design of synthetic PPIs but that *in vitro* derived PWMs provide a good starting point.

### Optimization of PPI is possible through additivity of substitution effect

In the PPI assay with double mutants, we found that substitutions generally have additive effects on PPI signal. Thus, we hypothesised that we could optimize PPI scores by combining more than two substitutions in the PWM_max_ backgrounds. In order to identify systematically binding preferences at each peptide position, we added all of the PWM_max_ peptide single mutants to the 3 PDZ peptides double mutant libraries described above (Figure 2A). In a second DHFR PCA bulk competition assay, we characterized the binding strength of the single and double mutants of the 3 peptide families to their designed PDZ domains, resulting in an additional 480 PBD-peptide variant pairs to the 1200 peptide variants from the double mutant libraries (Figure 2A). This assay was performed and analysed as above, except that we added a filter to remove outliers (see methods, Figure S6-S8).

**Figure 2.**
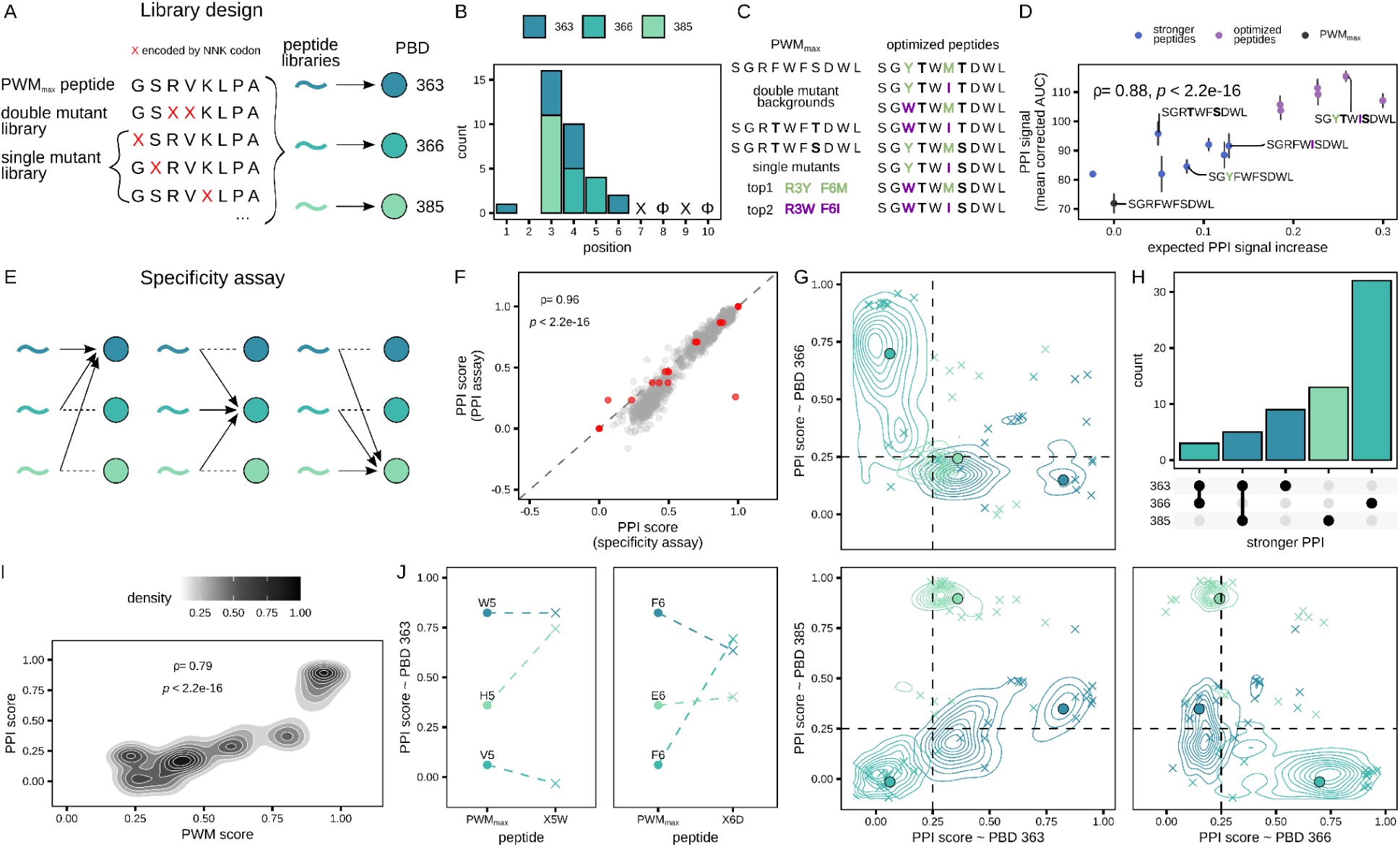
Profiling the specificity between peptide libraries and binding domains. **A.** Schematic of the library design strategy combining the double and single peptide mutants. Each peptide mutant is designed from the PWM_max_ peptide sequence of one PBD. **B.** Bar plot showing the number of single mutants per position in the peptide leading to a gain in PPI strength to its designed PBD. The core motif of PDZ domains (class II) is labelled on the plot, where X is any amino acid and Φ represents hydrophobic residues. **C.** Representation of the design approach to optimize peptide sequences. Starting from the PWM_max_ peptide sequence, two double mutant backgrounds were selected and two positions of single mutants. For each single mutant position, two amino acid substitutions were selected. Both double mutant backgrounds were combined with the all combinations of single mutants to result in eight PCA optimized sequences for each peptide family. **D.** Comparison of the mean PPI signal obtained from individual characterization (n=3) with the expected gain in PPI strength based on PCA bulk competition experiment of the 363 peptide family (see methods). Error bars show the standard deviation between replicates. Peptide sequences and the key mutated positions are labelled and colored according to the mutation rank. Spearman’s correlation coefficient and its p-value are labelled. **E.** Schematic illustrating the experimental design of the competition between peptide libraries to characterize the specificity of three PBDs, where the PPI strength of the three peptide libraries is assayed for each PBD. **F.** Scatter plot representing the correlation between the PCA scores obtained from PBD-peptide pairs present in the PPI assay and in the specificity assay. Red data points represent reference sequences. Spearman’s correlation coefficient and its p-value are labelled. **G.** Density plots comparing the PPI score of peptide variants against the three PBDs. The colors of the ellipses represent the peptide families. The circled data points represent the PWM_max_ peptides and the x data points are the selected variants for small-scale validation. The dashed lines highlight the quadrant where specificity of PBDs overlaps (top right). **H.** Upset plot shows intersections of the PBDs and the bar plot shows the number of peptide variants, single and double mutants, leading to a gain of interaction to these PBDs in comparison to their respective PWM_max_ peptide. The colors of the bars represent the peptide family of the peptide variant. **I.** Density plot showing the correlation between the PPI score obtained for each PBD-peptide pair and their PWM score. Density is represented by the shade of the ellipses. Spearman’s correlation coefficient and its p-value are labelled. **J.** Examples of single mutants from the 366 and 385 peptide families which can lead to a gain of interaction to PBD 363. The mutated residues from each PWM_max_ peptide are labelled above each point and the x are the mutation PPI scores. Colors represent peptide families.

From the single mutant libraries, we found that the positions of the substitutions leading to PPI gains are all upstream of the core binding motif, which supports the evidence that PDZ domain binding can be enhanced by modifying flanking residues (Zarin & Lehner, 2024) (Figure 2B). We hypothesized that amino acid properties allowing stronger binding *in vivo* would be different than the ones encoded in PWM_max_ peptides since the cell environment imposes different constraints than the *in vitro* settings in which the PWM_max_ peptides were defined. Thus, we compared the amino acid size and hydrophobicity between the PWM_max_ peptides and peptide variants leading to stronger PPIs. We found that PWM_max_ peptides are enriched in large residues and that peptide variants are less hydrophobic than the PWM_max_ peptides (Figure S13). This observation is surprising since we would expect that an increase in hydrophobicity strengthens the interaction. Indeed, the binding between peptides and PDZ domains is driven by hydrophobicity between the core motif and the binding site (Tonikian et al., 2008). However, it was shown that phage-display also enriches peptide variants towards higher hydrophobicity (Luck & Travé, 2011).

With many double and all single mutant effects characterized on PPI signal, we have the necessary information to test if substitution additivity applies to more than two mutations. We selected two double mutant backgrounds which showed significantly stronger PPI scores than the PWM_max_ peptides. Then, we selected two positions in the single mutant libraries of each PWM_max_ peptide at which the PPI scores were the highest. At these positions, we selected the top two scoring amino acids. By combining the two double mutant backgrounds with the 4 single substitutions, we obtained eight PCA optimised peptides for each PBD (Figure 2C). The F[3]-optimised peptide and the F[1,2]-PBD were encoded at an inducible locus in order to control the expression level and avoid reaching the DHFR PCA signal saturation (Aranda-Díaz et al., 2017; Lemieux et al., 2024). We performed growth curve measurements to characterise the optimised peptides for PPI strength using the inducible system.

We found that most of the optimised peptides gain PPI signal with their designed PBD (Figure S14A). To test if additivity is observed also at a higher order than with double substitutions, we compared the PPI signal obtained for the optimised peptides with the expected increase in PPI signal. The expected increase in PPI signal is computed by summing the difference in PPI scores obtained for single mutants compared to the PWM_max_ peptide in the bulk competition assay, which assumes perfect additivity. For the 363 peptide family, we observed that PPI signal can be optimised with predictability based on substitution additivity (Spearman’s correlation, ρ = 0.88, p < 2.2e-16, Figure 2D). However, for peptide families 366 and 385, the optimization is not as predictable (Figure S14C). Some substitutions, when combined, lead to decrease in PPI strength even if they showed increase in PPI scores when isolated (not all were significant). Despite that additivity is not generalizable for more than two substitutions, we were able to design a peptide sequence for each PBDs that showed stronger PPI than the single mutants or double mutant background (Figure S14C).

We have successfully increased PPI signal using additivity of more than three substitutions for the three PDZ domains selected. However, these results reveal that substitution additivity in PPIs is not universal and it can not always be used to optimize the interaction. Thus, multiple peptide sequences should be designed to avoid incompatible substitutions for strong PPIs.

### Peptide substitutions do not have the same effects on binding among PDZ domains

Above, we characterized the binding preferences of 7 PBDs and we observed differences from *in vitro* expectations, including stronger PPIs. However, stronger interactions could lead to less specific binding (Bendel et al., 2024), which would weaken the interest of these PBD-peptide pairs for synthetic system design. To test this possibility, we measured the binding specificity of the 3 PDZ domains (PBD 363, PBD 366 and PBD 385) to each of their peptide families including all single and double mutants described previously, resulting in > 5000 PBD-peptide variant pairs (Figure 2E). We defined a change in binding specificity of peptide variants as a significant PPI signal increase relative to their PWM_max_ peptide, i.e. when a peptide variant has a stronger PCA signal to a non-designed PBD than its PWM_max_ peptide background. This assay was performed in the second DHFR PCA bulk competitions with the single mutant libraries and analysed in the same way (see methods, Figure S6-S8). We compared the PPI scores obtained from the PPI assay and from the specificity assay since many PBD-peptide variant pairs were present in both experiments. We observed a strong correlation between the two assays (Figure 2F, Spearman’s correlation, ρ = 0.96, p-value < 2.2e-16), indicating great replicability. Individual validations were performed for 150 PBD-peptide pairs and we obtained consistent results between the PCA competition and the small-scale assay (Spearman’s correlation, ρ = 0.77 to 0.92, p-value < 6.2e-7, Figure S9C).

By characterizing the PPI strength of three peptide families against all three PBDs, we assayed the effect of single and double mutants on the binding specificity from different background peptides. We observed that most peptide variants bind specifically to the PBD used to define the peptide families (Figure 2G, Figures S12-14). However, we detected a significant change in binding specificity for 154 peptide variants from one PBD to another relative to their family PWM_max_ peptide (Figure 2H). Most of these peptide variants are part of the 363 and 385 peptide families and we observe that their corresponding PBDs have overlapping binding preferences. The large number of shared binders between PBDs is consistent with the data from the PWM, since they share three essential amino acids for binding (S7, W9, L10, Figure S1) that are encoded in their PWM_max_ peptide.

However, we observed that peptide variants leading to a gain of PPI signal to their designed PBD compared to their PWM_max_ peptide, do so mostly in a specific manner (Figure 2H). A minority (8/62) of peptide variants which increase binding also lead to a gain of PPI signal to another PBD, indicating that substitutions increasing binding do not affect all PDZ-peptide interactions in the same manner. Since we tested the PPI of the PDZ domains with a much higher peptide diversity than in the previous experiment by adding all the single mutants and testing specificity between PBDs, we postulated that the PWMs would be able to better predict *in vivo* binding. We compared the specificity characterization obtained by DHFR PCA and by computing a PWM score of each peptide variant with the PWM of the three PBDs tested. We observe that the PWM scores and the PPI scores are correlated significantly (Figure 2I, Spearman’s correlation coefficient = 0.79, p-value <2.2e-16). The relationship is much stronger for the libraries used to test specificity between PBDs (Figure 2A & E), which cover a much broader range of PWM scores, than for the double mutant libraries (Figure 1I, Figure S5). Combined with the observation that amino acid properties leading to the strongest PPIs *in vivo* are not the same as the ones encoded in the PWM_max_ peptide, this finding indicates that a PWM is a powerful tool to predict non-binding peptides to a PBD, but it has limited use to find the best binding peptide to a PBD *in vivo*.

Since we found that amino acid properties are different between stronger binding peptides *in vivo* and PWM_max_ peptides, we asked if the same substitutions in different peptide backgrounds lead to the same effect on binding. Looking at specific examples of the three peptide families binding to PBD 363, we found that X5W leads to a gain of PPI when introduced in the 385 PWM_max_ peptide but has a slightly negative effect on binding in the 366 PWM_max_ background (Figure 2J, left). We observed another case where both peptide backgrounds have a F6D with opposite effects. In the 363 PWM_max_ background, F6D led to a decrease in binding but, in the 366 PWM_max_ background, it led to a gain of PPI (Figure 2J, right). These examples show that single substitutions in a peptide do not affect their binding to the same PBDs in a singular way and that the peptide background sequence is critical to define substitution effect on the PPI. Thus, substitution effects within one peptide family do not transfer across different peptide backgrounds even if a lower amino acid hydrophobicity and size are favorable for binding in all backgrounds (Figure S13).

Taken together, our results reveal that a gain in PPI signal to one PBD does not equate to a gain to all PBDs of its family. For peptide variants leading to stronger PPIs than the PWM_max_ peptides, we observed that their gain is very specific to their designed PBDs and that this is mostly driven by lower peptide hydrophobicity and amino acid size outside of core motif positions. It remains to be investigated whether the binding strength between these peptide variants and their PBD is increased by a change in affinity, or whether another mechanism explains the observed gain in PPI signal.

### Binding energies do not explain PPI signal gains

To determine whether peptide variants with stronger PPI signals also form more stable complexes, we modelled the interactions between the three PDZ domains and their corresponding PWM_max_ peptides. Complexes were predicted with Alphafold3 (n = 5) (Abramson et al., 2024) and validated against available experimental structures of the same domains. The predicted models showed excellent agreement with their crystallographic counterparts, with backbone Cα RMSDs of 0.81, 0.71 and 0.50 Å for PBD 363, PBD 366 and PBD 385 respectively. For PBD 363 and PBD 385, whose crystal structures contain peptide ligands sufficiently similar to the modelled sequences, peptide conformations were also accurately reproduced (Cα RMSD = 0.17 and 0.12 Å, respectively).

The models revealed distinct peptide-binding modes among the three PDZ domains (Figure 3A). The PWM_max_ peptide bound to PBD 366 adopts an extended β-sheet conformation along the binding groove, resulting in extensive contacts across the peptide sequence and explaining why substitutions at most positions influence PPI strength (Figure S11). In contrast, the PWM_max_ peptide bound to PBD 385 interacts primarily through its five C-terminal residues, whereas the N-terminal half remains conformationally heterogeneous across models. Consistent with this observation, mutations within the N-terminal region have little effect on PPI strength (Figure S12). These structural differences provide a plausible explanation for the distinct mutational sensitivities and binding specificities observed among the PDZ domains. We next asked whether the stronger PPI signals detected by DHFR PCA could be explained by improved binding energetics. Changes in binding energies (ΔΔG) were calculated for all single and double mutants using FoldX (Schymkowitz et al., 2005) and RosettaFlexddG (Barlow et al., 2018; Chaudhury et al., 2010). The two approaches showed substantial agreement overall (Spearman’s correlation, ρ=0.77, p-value < 2.2e-16, Figure 3B), particularly for substitutions predicted to have modest energetic effects (ΔΔG < 1 kcal/mol). Comparison with experimental PPI scores revealed significant correlations for two of the three PDZ-peptide systems (Figure 3C), indicating that computational estimates capture part of the variation observed *in vivo*.

**Figure 3.**
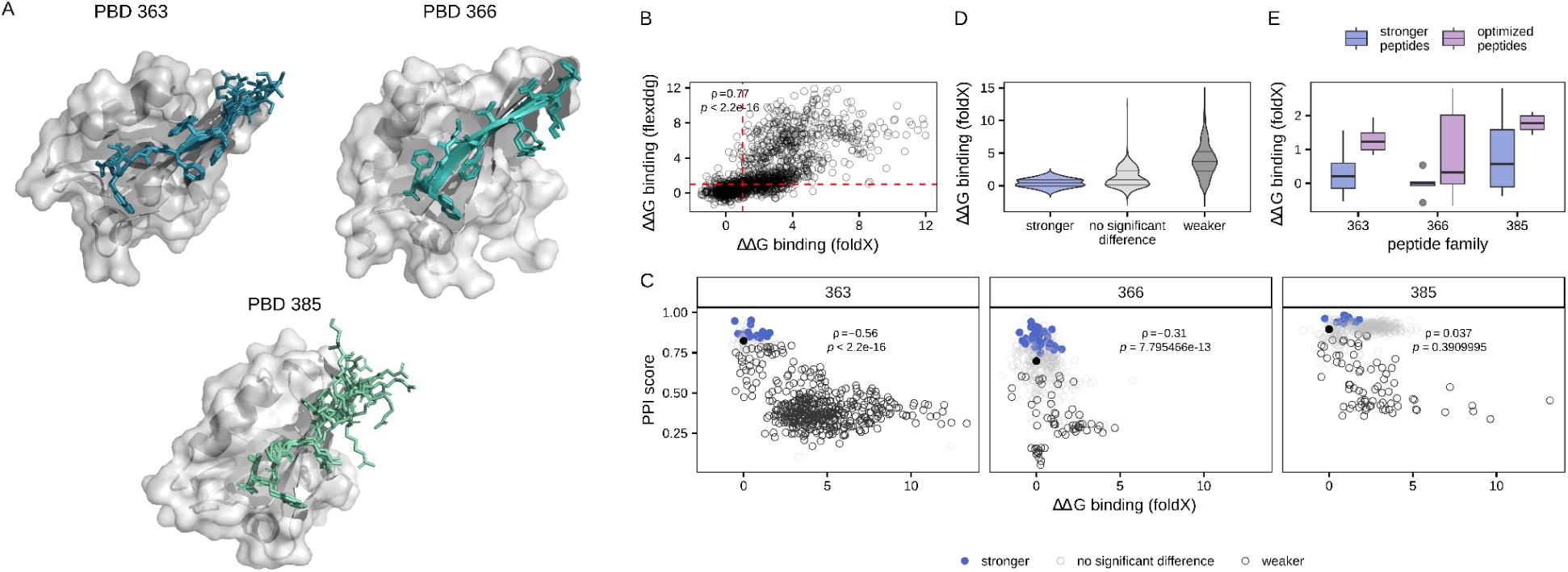
*In silico* modelling and energetic analysis of PDZ-peptide interaction. **A.** AlphaFold3 models of PBD 363, PBD 366 and PBD 385 bound to their respective PWM_max_ peptides. Peptides are shown in color and the five predicted models are superposed on the receptor structure. **B.** Comparison of ΔΔG values predicted by FoldX and RosettaFlexddG for single and double mutants. The dashed red lines indicate ΔΔG=1. Scatter plots represent individual peptide variants and the Spearman correlation coefficient and associated p-value are labelled. **C.** Relationship between predicted ΔΔG values and PPI scores for each PBD-peptide family. Colors illustrate the experimental classification of peptide variants relative to the PWM_max_ peptide based on DHFR PCA. Spearman’s correlation coefficients and p-values are labelled. **D.** Violin plot showing the distributions of FoldX-predicted ΔΔG values for peptide variants classified as stronger, neutral (no significant difference) or weaker binders by DHFR PCA. Positive ΔΔG values indicate mutations predicted to destabilize the interaction relative to the PWM_max_ peptide. **E.** Distribution of FoldX-predicted ΔΔG values for peptide variants displaying stronger PPI signals than their PWM_max_ peptide and for the PCA-optimized peptides. Boxes show the median and interquartile range, with individual variants represented as points.

Despite these significant correlations, binding-energy predictions failed to explain most gains in PPI signal. Peptide variants classified experimentally as stronger binders frequently displayed positive ΔΔG values, indicating predicted destabilization of the interaction (Figure 3D). Among the 62 peptide variants that increased PPI signal relative to their PWM_max_ peptide, only 16 were predicted by FoldX to stabilize the complex (ΔΔG < 0). This overlap remained greater than expected by chance (Fisher’s exact test, p-value = 0.00258), revealing that predictions do capture some of the stronger binding but only marginally. Similar results were obtained with RosettaFlexddG.

Finally, we examined whether the additive effects observed experimentally could be explained by cumulative energetic stabilization. The optimized peptides generated by combining beneficial substitutions were predicted to be largely destabilizing according to both FoldX and RosettaFlexddG (Figure 3E), including variants from the 363 peptide family that displayed the largest increases in PPI signal (Figure 2J). Together, these results indicate that structural models capture the architecture of PDZ-peptide interactions and partially reflect mutational effects gains observed *in vivo.* However, changes in binding energy alone cannot account for the majority of PPI signal accurately, suggesting that additional cellular factors contribute to the enhanced interaction signals detected by DHFR PCA.

### Stronger PPI signal stems from higher availability of peptide variants

An increased peptide abundance without a gain in binding energies could be an alternative explanation for the PPI signal gains observed with DHFR PCA since this method relies on *in vivo* binding events. We tested this hypothesis using a DHFR PCA experiment that can detect changes in protein abundance. In a yeast strain expressing from its genome a YFP-F[1,2] fusion protein (Figure 4A) (Levy et al., 2014), we expressed the F[3]-peptide libraries without a PBD. The binding events between the F[3]-peptide and the YFP-F[1,2] lead to the mDHFR complementation that strongly correlates with the abundance of both proteins (Levy et al., 2014). Since YFP-F[1,2] abundance is constant and the peptide sequences are variable, the PCA signal detected reflects the abundance of the F[3]-peptide variants and thus, their availability for binding to the PBD in the cytosol in the previous assays described above. This availability would be limited for instance by degradation or by sequestration with yeast endogenous proteins. We performed bulk-competition experiments to characterize binding availability of all the peptide variants characterized for PPI. PCA scores were computed using reference sequences as described above. We obtained high replicability between triplicates (ρ > 0.91, p-value < 2.2e-16, Figure 4B) and we performed small-scale validations which showed good reproducibility (Figure 4C, Spearman’s correlation, ρ = 0.88, p-value = 0.002, Figure S9D).

**Figure 4.**
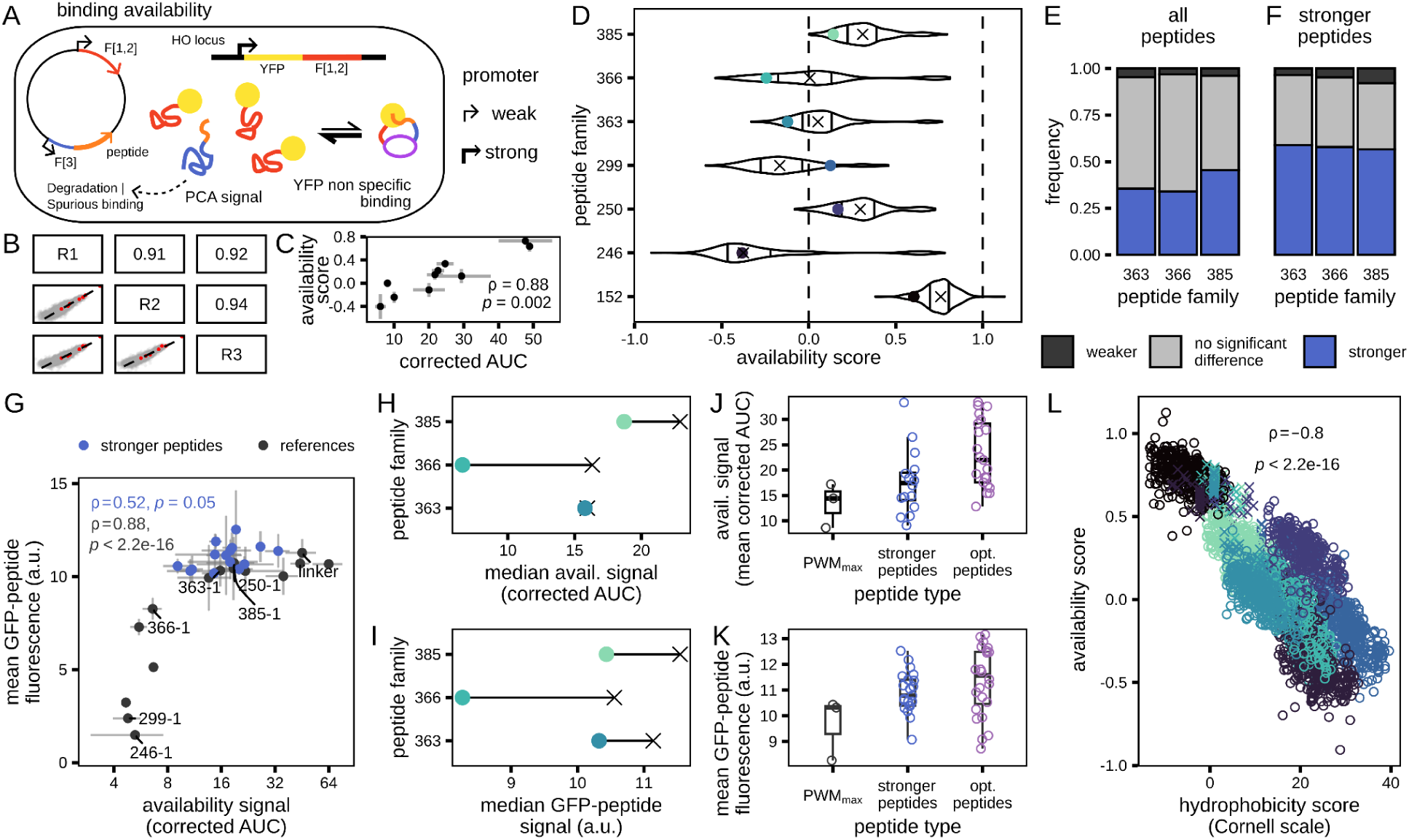
Effect of mutation on peptide binding availability and abundance. **A.** Schematic of the PCA binding availability assay. A YFP-F[1,2] is highly expressed from the genome and the F[3]-peptide libraries are expressed from a plasmid vector. PCA signal is detected when the F[3]-peptide and YFP-F[1,2] bind non specifically (Levy et al., 2014). **B.** Scatter plots representing the correlation between the normalised *s* from each replicate. The replicates are identified on the diagonal and the Spearman’s correlation coefficients are shown on the mirror quadrant to each scatter plot (p-values < 2.2e-16 for all comparisons). The red data points correspond to the reference sequences added in the libraries. **C.** Scatter plot showing the comparison between PPI scores and the PCA signal obtained from individual growth curve analysis (corrected AUC). Error bars represent the standard deviation obtained for each experiment performed in triplicate. The Spearman’s correlation coefficient and its p-value are labelled. **D.** Violin plot representing the PPI score distributions for each peptide family. The colored data points are the PWM_max_ peptides. The median of each distribution is shown as a cross and the 25th and 75th quartiles are drawn as lines in the violins. The dashed lines at 0 and 1 represent the sensitivity range of the assay covered by the reference sequences. **E.** Bar plot representing the peptide variants that showed differences in availability compared to the PWM_max_ peptide of their peptide family and **F.** only for the peptide variants that showed increased PPI score compared to their PWM_max_ peptides. **G.** Scatter plot comparing the availability signal obtained from individual characterization of peptide sequence availability (n = 3) with the fluorescence signal (arbitrary units – a.u.) of GFP-peptide (n = 3). The error bars show the standard deviation between replicates and the data point color indicates the peptide variants with stronger PPI score than their PWM_max_ peptides. The annotated data points are the PWM_max_ peptides and the positive control (linker). Spearman’s correlation coefficients and their p-values are labelled. **H.** Dumbbell plot showing the difference between the median availability signal (corrected AUC) and **I.** GFP-peptide fluorescence signal (a.u.) of the PWM_max_ peptides (point) and the stronger interacting peptides (cross). **J.** Distributions of the mean availability signal (corrected AUC) and **K.** GFP-peptide fluorescence signal (a.u.) obtained from individual characterization (n=3) of the PWM_max_ peptide, stronger interacting peptides and PCA-optimized peptides. **L.** Comparison between peptide availability and hydrophobicity scores obtained by summing the hydrophobicity of amino acids present in the peptide variants. The color of the data point shows the peptide family (same as in D) and the cross show peptide variants shortened by non-sense mutations. Spearman’s correlation coefficient and its p-value are labelled.

We observed that the PWM_max_ peptides of 6/7 peptide families have availability scores below their family median indicating that most peptide variants lead to an increase in binding availability (Figure 4D, Figure S15, S17A). We hypothesised that the presence of degron motifs, which are sequences known to favor protein degradation, in the PWM_max_ peptides could explain their low availability. Thus, we searched for known yeast degron sequences in the PWM_max_ peptides and in all peptide variants, using the Degronopedia database (Szulc et al., 2024). We found that 467/3466 of the characterized peptides variants contained a degron motif (Figure S16). We would expect C-terminal degrons to impact the most the assay since all peptide variants are present in C-terminal of the F[3], but none were C-terminal degron. Most of the degron found (400/467) are present in the 246 peptide family because the internal degron RLLL is present in all variants, which explains that the whole peptide family has a low availability. The other degron sequences were found in peptide variants of the 363, 366 and 385 families, respectively 22, 2, and 1 instances. Internal degrons were found (20/22) in the peptide family 363 and these peptide variants (14/20) have generally a lower availability than the family median. The remaining degron sequences were N-terminal degron and, as expected, their effect on availability is limited since 3/5 show availability higher than the 25th percentile of their family distribution. Globally, the presence of degron sequences explains part of the peptide availability, but most of the peptides do not encode known degrons. Knowing that most peptide variants do not encode a degron sequence, we then asked if increased availability led to an increase in PPI signal by comparing the PPI and availability DHFR PCA results. Looking at the PDZ peptide families, the global proportion of peptide variants with significantly higher availability than the PWM_max_ peptide is 38 % (Figure 4E), while, when considering only the stronger binding peptides, the proportion raises to 57 % (Figure 4F). This observation suggests that an increased binding availability can contribute to increased PPI signal.

Since active degradation does not explain most of the availability variance, we postulated that a more subtle variation in abundance could contribute to the availability signal and explain PPI signal gains. Thus, we tested if the increased availability resulted from increased abundance by fusing peptide variants to the C-terminal of the GFP and measuring cell fluorescence by flow cytometry. First, we verified that the fluorescence measurements obtained from GFP-peptide were replicating our results by selecting reference sequences which showed a wide range of availability signal (Figure 4G). We detected a strong correlation between the two assays (Spearman’s correlation, ρ = 0.88, p < 2.2e-16), although we observed a saturation of the fluorescence signal as availability measurements increased. The protein stability fused to the peptide could explain the different range of sensitivity, since the GFP has a stable fold and could stabilize abundance for some peptides which have a lower abundance when fused to F[3] (Ho et al., 2018) which is expected to be disordered (Pelletier et al., 1998; Remy & Michnick, 2007). Doing the same measurements with peptide variants which showed stronger PPIs (Figure 4G), we found that their median availability is higher than the PWM_max_ peptide for each peptide family (Figure 4H). Consistently, the median fluorescence signal of GFP-peptide variants tested is higher than the PWM_max_ peptides (Figure 4I). We also observed a general increase in availability signal (Figure 4J, Figure S14B) and in GFP-peptide fluorescence signal (Figure 4K) for the optimized peptides compared to the PWM_max_ peptides and to the peptide variants which showed stronger PPIs.

Since stronger binding peptides were enriched in less hydrophobic residues and that PPI optimization is associated with increased peptide availability *in vivo* (Figure S13), we postulated that peptide hydrophobicity could be a predictive sequence feature for availability. Thus, we compared the hydrophobicity score (Cornell scale) (Karplus, 1997) and the availability scores of each peptide variant and found that they are strongly correlated (Figure 4L, Spearman’s correlation, ρ = –0.8, p < 2.2e-16). This relationship is particularly strong when looking at shortened PWM_max_ peptides by premature stop codons. The longer the peptides are, the more hydrophobic they become and the lower the availability is (Figure S17B). Since each PWM_max_ peptide variant has a different starting hydrophobicity that we alter slightly by mutating one or two residues, combining them to look at the overall correlation between availability and hydrophobicity could be artefactual. However, the relationship between availability and hydrophobicity is still significant when we isolate each peptide family and remove the stop codons from the comparison (Figure S17C, Spearman’s correlation, ρ = –0.49 to –0.77, p < 2.2e-16). This finding suggests that we can use amino acid hydrophobicity to guide peptide design towards higher availability. Because peptides with stronger PPIs show higher availability, the use of peptide hydrophobicity as a feature could help to design stronger binders while preserving key residues for binding in synthetic PPIs *in vivo*.

We found that peptides which are very hydrophobic are less available, without encoding for known degron sequences. However, the wide range of availability signal is not explained by degradation and it suggests that hydrophobic peptides could have more spurious interactions with endogenous protein than less hydrophobic peptides and lead to their sequestration. To explore this hypothesis, we performed a simple analysis comparing the relative hydrophobicity of the YFP-F[1,2] to the rest of the yeast proteome showing that the YFP-F[1,2] is less hydrophobic than the average yeast protein (Figure S18). Thus, the negative relationship between availability and hydrophobicity could be explained by the higher frequency of spurious interaction of hydrophobic peptides with the more hydrophobic proteins present in the cytosol, limiting the availability of the peptide to bind with the YFP-F[1,2].

Taken together, we found evidence that peptide variants which gain PPI strength *in vivo* are enriched in peptide sequences that have an increased availability compared to the PWM_max_ peptides. The rational design of sequences to optimize availability is easily achievable since availability is defined by peptide hydrophobicity. Combined with the known binding preferences of PBDs, adding availability as a design feature would increase binding signal *in vivo* for synthetic PPIs of interest in biotechnology and therapeutics development.

## Discussion

In this study, we investigated the extent to which binding data derived from *in vitro* experiments predict how PBDs and peptides assemble in living cells. We found that the information contained in PWMs derived from phage-display is very informative to design a background peptide sequence binding to PBD which can then be mutated to enhance PPI signal *in vivo* (Benz et al., 2025). Using DHFR PCA, we detected a strong additive effect on PPI signal from double mutants of the PWM_max_ peptides and this effect can be extended up to four substitutions. Additivity of mutation effects in peptide binding to PDZ domains was demonstrated using affinity-based assays (Stiffler et al., 2007; Tahti et al., 2023). However, since PWMs were not able to predict the increased interaction signal of combined substitutions, we reasoned that the additive effects do not stem solely from increased affinity. Then, we answered two questions addressing PDZ domain specificity. First, we asked whether peptide increasing binding led to decreased specificity to the canonical PDZ. Second, we asked if the PDZ domain specificity *in vivo* was the same as characterized *in vitro*.

We found that the peptides increasing PPI signal were very specific to one PDZ. This finding does not agree with the original classification of PDZ domains, since it was postulated that PDZ domains from the same class (I, II and III) would bind non-specifically the same peptide as long as it encodes the class core motif (Amacher et al., 2020). Indeed, in our assays, we use PBD 363 and 385 which share their core motif (S/T-X-W-L) but the strongest peptide binders also display specificity since binding relies as much on the residues outside the core motif as those within (Amacher et al., 2026). The classification of PDZ domains has been criticized for more than a decade, but the systematic characterization of specificity using the same peptide variants *in vivo* against multiple PDZ domains was lacking. Our work contributes to the evidence that PDZ specificity is a continuum and that PDZ selectivity *in vivo* is possible (Amacher et al., 2020; Stiffler et al., 2007).

Our structural models are consistent with this interpretation. Although PBD 363, PBD 366 and PBD 385 belong to the same PDZ class, AlphaFold3 predicted distinct peptide-binding modes for each domain. In particular, the PWM_max_ peptide bound to PBD 366 formed an extended β-sheet along the binding groove, whereas binding to PBD 385 relied primarily on interactions involving the five C-terminal residues. These differences provide a structural explanation for the distinct mutational sensitivities observed experimentally and further support the conclusion that residues outside the canonical PDZ motif contribute substantially to binding specificity. This observation is consistent with previous studies showing that non-core residues can modulate both affinity and selectivity in PDZ-peptide interactions (Amacher et al., 2026; Tahti et al., 2023). Consistent with previous affinity-based measurements, our structural models further suggest that the energetic contribution of individual peptide residues depends on the extent of their contacts with the PDZ binding groove. The nearly continuous interaction surface observed for PBD 366 may explain why substitutions across most positions influence PPI strength, whereas the more localized interactions predicted for PBD 385 are compatible with the tolerance of its N-terminal residues to mutation.

We showed that the overall PDZ domain specificity characterized *in vitro* is conserved when studying their interaction in living cells. This question is highly relevant since most PDZ-peptide interactions are still characterized using *in vitro* approaches. PDZ domains are involved in a wide range of cellular processes and repurposed in cancer and virus infections (Mészáros et al., 2021; Mihalič et al., 2023; Rrustemi et al., 2024). Promising therapeutic treatments design PDZ inhibitors which are often specific competitive peptide inhibitors. The limiting step to design peptide inhibitors is often the function in the cell host due to the cross-selectivity with endogenous PDZ domains (Nardella et al., 2021). Thus, better understanding how synthetic peptides behave and interact in cells is essential and there is a need to understand how *in vitro* specificity assay transfers to *in vivo* settings. Our results show that specificity is conserved for three human PDZ domains in *S. cerevisiae*, but it might not be the case for other host cells. Indeed, yeast is a frequently used model organism to characterize protein function and model diseases from more complex organisms due to the conservation of proteostasis and organelle function with higher eukaryotes (Do et al., 2026; Kachroo et al., 2022). However, it is unknown how our findings translate to human cells given the higher complexity of the human proteome and the additional layers of control over protein expression (Do et al., 2026).

Neither phage-display-derived PWMs nor structure-based ΔΔG predictions were sufficient to explain the largest gains in PPI signal observed *in vivo*. The energetic calculations nevertheless captured part of the experimental variation. Predicted ΔΔG values were significantly correlated with PPI scores for two of the three systems, and variants predicted to stabilize binding were enriched among peptides displaying stronger PPI signals. Thus, binding energetics contribute to the observed interaction strengths. However, most peptide variants exhibiting improved PPI signals were predicted to have neutral or destabilizing effects on binding, indicating that affinity changes alone cannot account for the strongest gains observed *in vivo*. This observation suggested that factors beyond binding affinity contribute substantially to interaction strength in cells. Among these, peptide abundance and availability appeared particularly plausible because DHFR PCA reports productive interactions between molecules present in the cellular environment. We therefore investigated whether peptide availability could contribute to the increased PPI signals observed for some variants. Peptide abundance is expected to be a key factor of availability because it directly affects the fraction of molecules accessible for binding. As demonstrated by Levy et al., 2014, the availability of a protein in living yeast cells can be assayed using DHFR PCA. Consistent with our expectations, we found an association between increased peptide availability and increased PPI signal between domains and peptides. This observation suggests that availability of a peptide is a key feature to consider when designing synthetic PPIs. However, availability is not a direct measurement of abundance. Thus, to determine if peptide abundance was the main driver of peptide availability, we measured the fluorescence intensity of cells expressing GFP-peptide constructs. We found a strong agreement between availability and fluorescence measurements. However, the sensitivities range of each assay were different and we observed signal saturation for the fluorescence assay while the availability values kept increasing. This observation could be related to the different effects of the two protein fusions used in both assays (Ho et al., 2018), but it could also be caused by the sequestration of peptide variants by endogenous proteins. This mechanism could also explain the signal saturation observed for fluorescence measurements while availability had a broader range of signal, since hydrophobic peptides would not change in abundance but only in their ability of diffusing freely. Indeed, spurious binding occurs also between peptide variants and endogenous proteins in the cell, as it can happen between endogenous proteins due to the noisy nature of interactomes (Levy et al., 2009). The hypothesis that reducing hydrophobicity of a peptide sequence could limit spurious interactions with the host proteome would require further experiments, but it is of significance for synthetic biology and therapeutics design.

Overall, our results suggest that the PPI signal reported by DHFR PCA is affected by cellular mechanisms that are absent from phage-display experiments or other *in vitro/in silico* assays. These effects contribute to modifying the peptide availability which in turn impacts the PPI signal obtained from a PBD-peptide pair. We found that peptide hydrophobicity is a very strong predictor of its availability and we propose that hydrophobicity could be used to alter peptide sequences for PPI engineering. The complexity of engineering living organisms, with their intricate machinery and interactomes, emphasizes the relevance of *in vivo* measurements to guide those modifications. Together, our results suggest that designing strong peptide-mediated interactions *in vivo* requires accounting for two distinct layers of information: sequence determinants that govern molecular recognition and sequence determinants that influence peptide availability within the cellular environment. Structural models, affinity measurements and energetic predictions capture important aspects of the former, but only partially account for the latter. Integrating both layers will be essential for the rational design of robust and specific synthetic protein-protein interactions in living cells.

## Author contributions

P.LE., A.K.D. and C.R.L. were involved in the experimental design. P.LE. and A.K.D. designed molecular biology material for this study. P.LE. performed the wet lab experiments. X.B. performed the *in silico* predictions. P.LE. S.D. and X.B. contributed to the data analysis. P.LE. wrote the first draft of the manuscript. P.LE., X.B., S.D., A.K.D., P. LA. and C.R.L. contributed to writing, revising and approving the final version of the manuscript. C.R.L. and P.LE. acquired the funds to support this project. P. LA. and C.R.L. supervised this work.

## Acknowledgments

We would like to thank Guillaume Diss who provided key materials and comments for the experimental design. We are thankful for the comments of Isabelle Gagnon-Arsenault on the manuscript, and for the support of all other members of the Landry Lab.

## Material availability statement

Raw sequencing data are deposited as an SRA BioProject (PRJNA1499819). All growth measurements, flow cytometry data and supplementary material including the code for data analysis and figure generation is available on the Github repository (https://github.com/Landrylab/Lemieux_et_al2026.git). The strains and plasmids are available upon request. Supplementary File 1 contains the Supplementary Tables 1 to 8.

## Supplementary Figures

**Figure S1.**
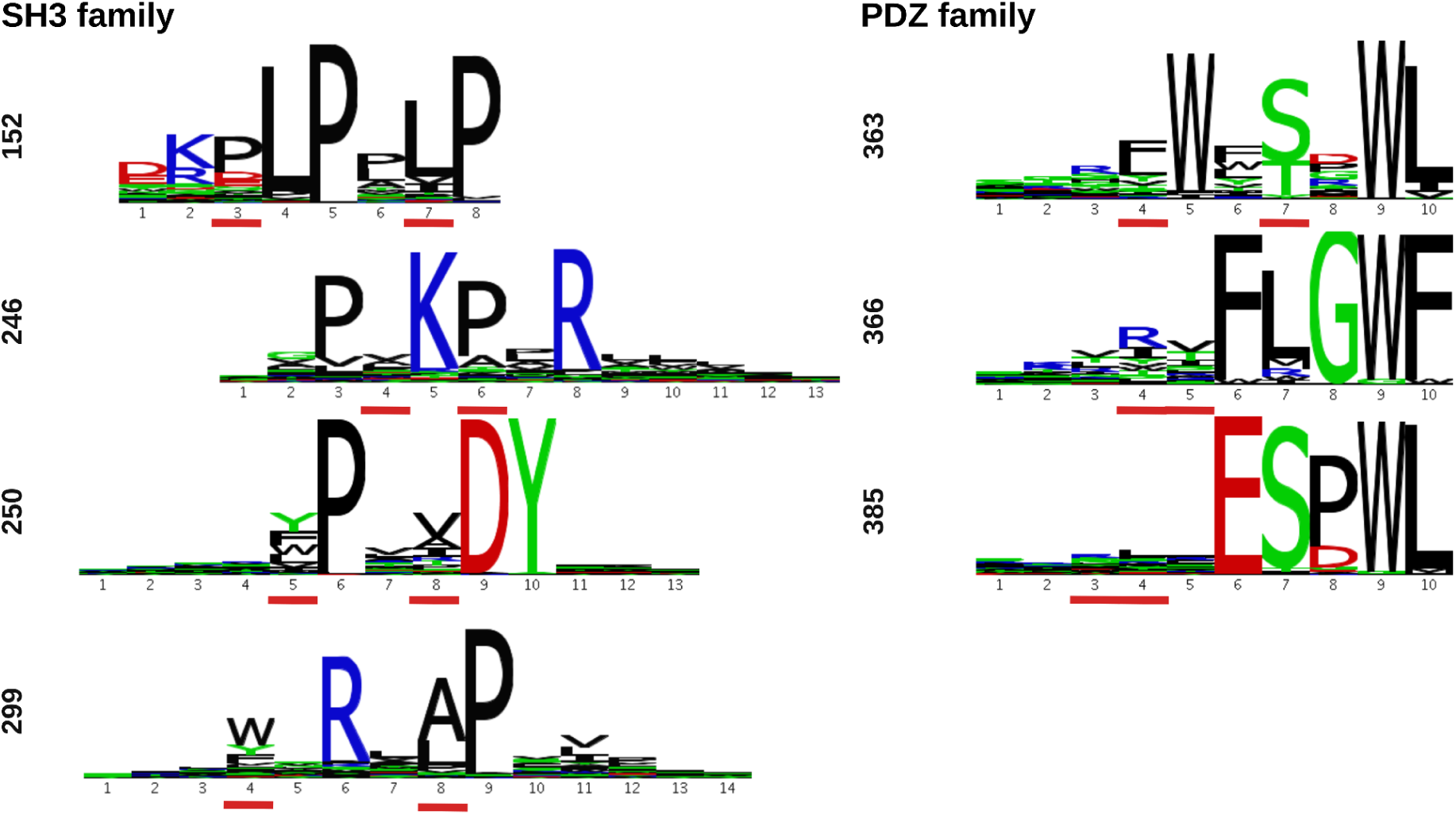
Specificity of the selected PBDs. The PWMs extracted from the PRM-DB (Teyra et al., 2020) are aligned to show the common binding motif of each PBD family. The underlined positions were degenerated with a NNK codon to create the mutant libraries for each peptide family.

**Figure S2.**
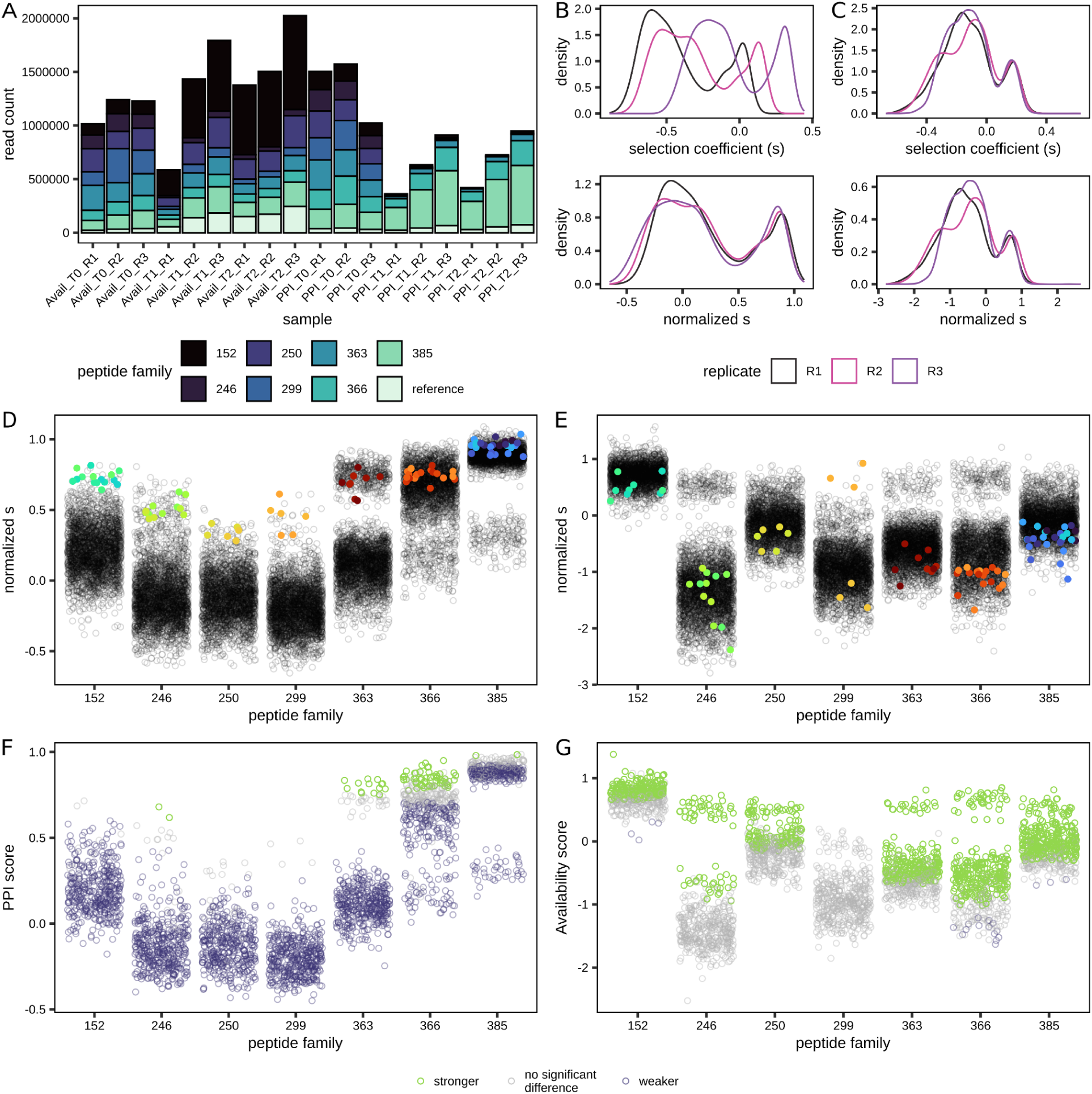
Processing of the sequencing results from the bulk competition assays performed to assess PPI strength and binding availability. A. Bar plot showing the read count of each sample where the colors represent the proportion of reads associated with each peptide family. The x-axis described each sample with the assay type, the time point and the replicate. B. Density plots illustrating the normalisation of the selection coefficient (see methods) computed between T0 and T2 for the PPI assay and C. the binding availability assay. D. Representation of the distribution of the PPI normalised s and E. the binding availability normalised s for each DNA sequence associated with their peptide family on the x-axis. The colored data points show the synonymous sequence encoding for each maximal binding peptide defined by the PWMs in each replicate. F. PPI and G. Availability scores are shown for each peptide sequence associated with their peptide family on the x-axis. The colors of the data point represent the classification of each peptide as a statically stronger (green) or weaker (violet) binder compared to the maximal binding peptide defined by the PWMs (Wilcoxon test, Benjamini-Hochberg correction, p-value < 0.05, see methods).

**Figure S3.**
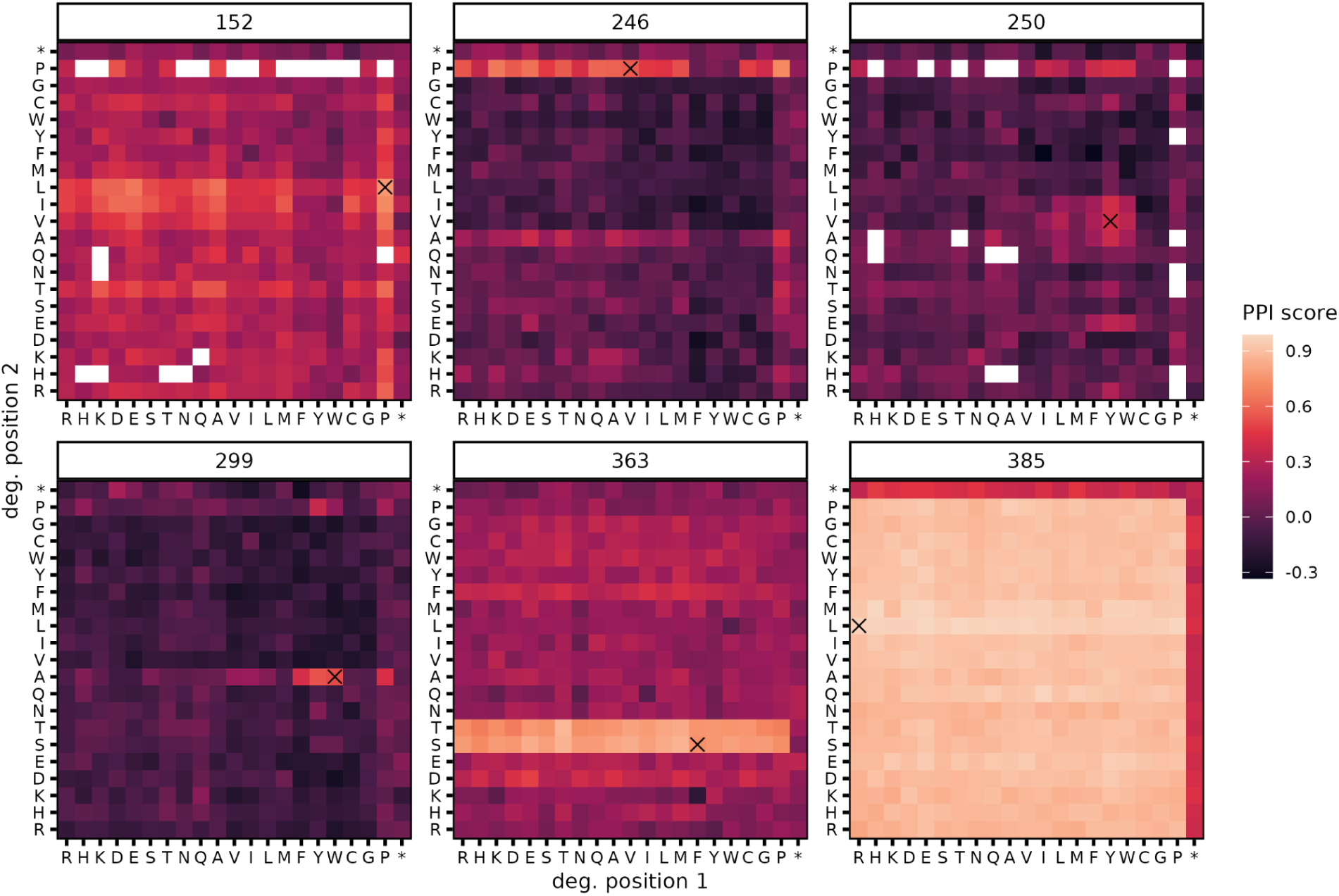
Relative PPI strength estimated from DHFR PCA in bulk competition. The paired PBD-peptide families are labelled on top of each panel. The color scale of the heatmaps represents the PPI scores and the pairs of mutated residues are labelled on the axis. Amino acid order follows their chemical properties (charged: R, H, K, D, E; polar: S, T, N, Q; hydrophobic: A, V, I, L, M, F, Y, W; special: C, G, P; stop: *). The cross highlights the PWM_max_ peptide sequence. White tiles are missing data.

**Figure S4.**
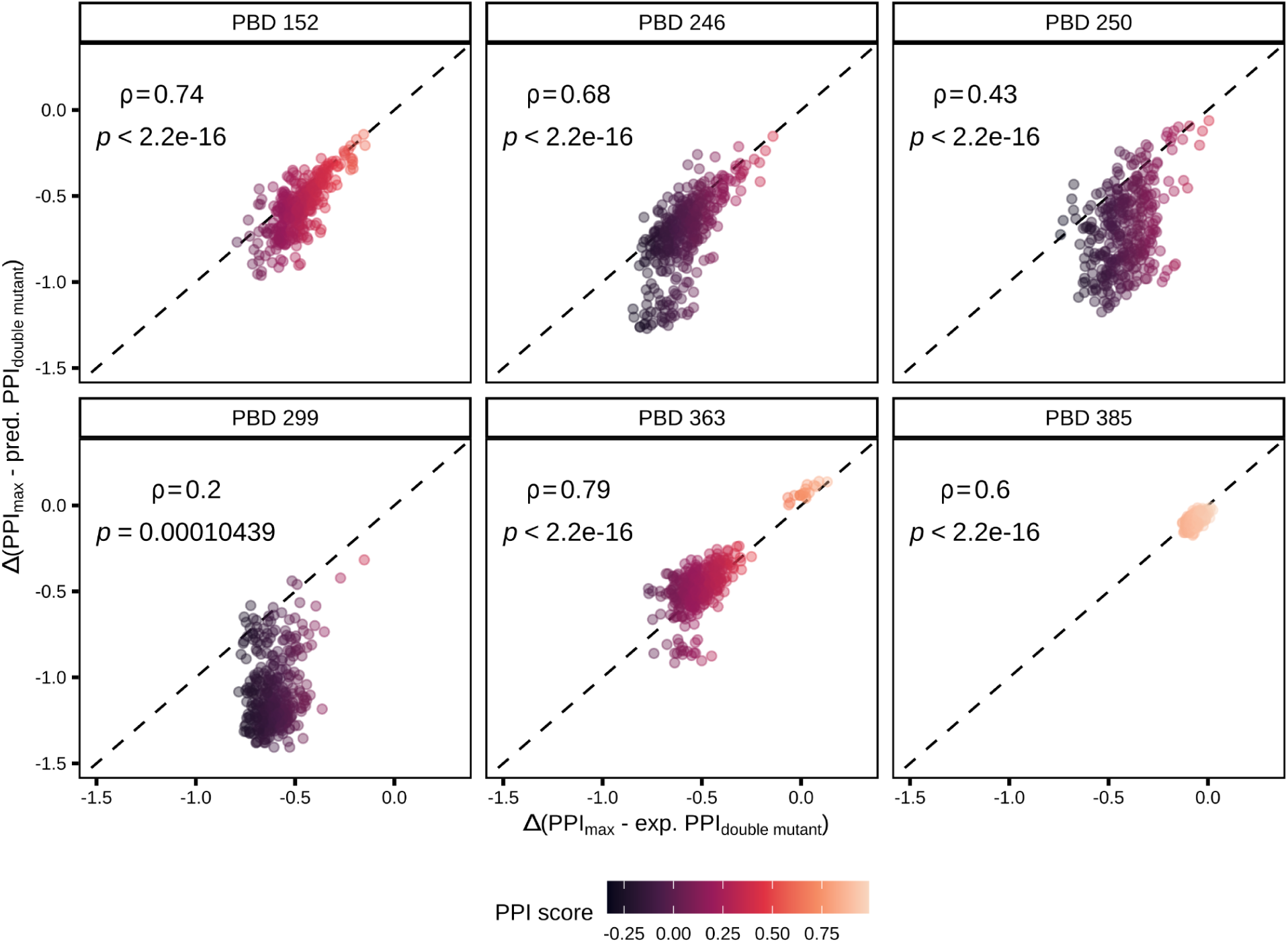
Expected versus observed PPI strength for double mutant peptides. Scatter plots showing the comparison between the difference in PPI score of the PWM_max_ peptide and the double mutants PPI score (exp. PPI_double mutant_, x-axis) and the difference in PPI score of the PWM_max_ peptide and the sum of PPI scores of single mutants (pred. PPI_double mutant_, y-axis). The PBDs are labelled above each plot. Spearman’s correlation coefficient and p-value are labelled for each comparison. The color of the data points indicates the PPI score of the double mutants.

**Figure S5.**
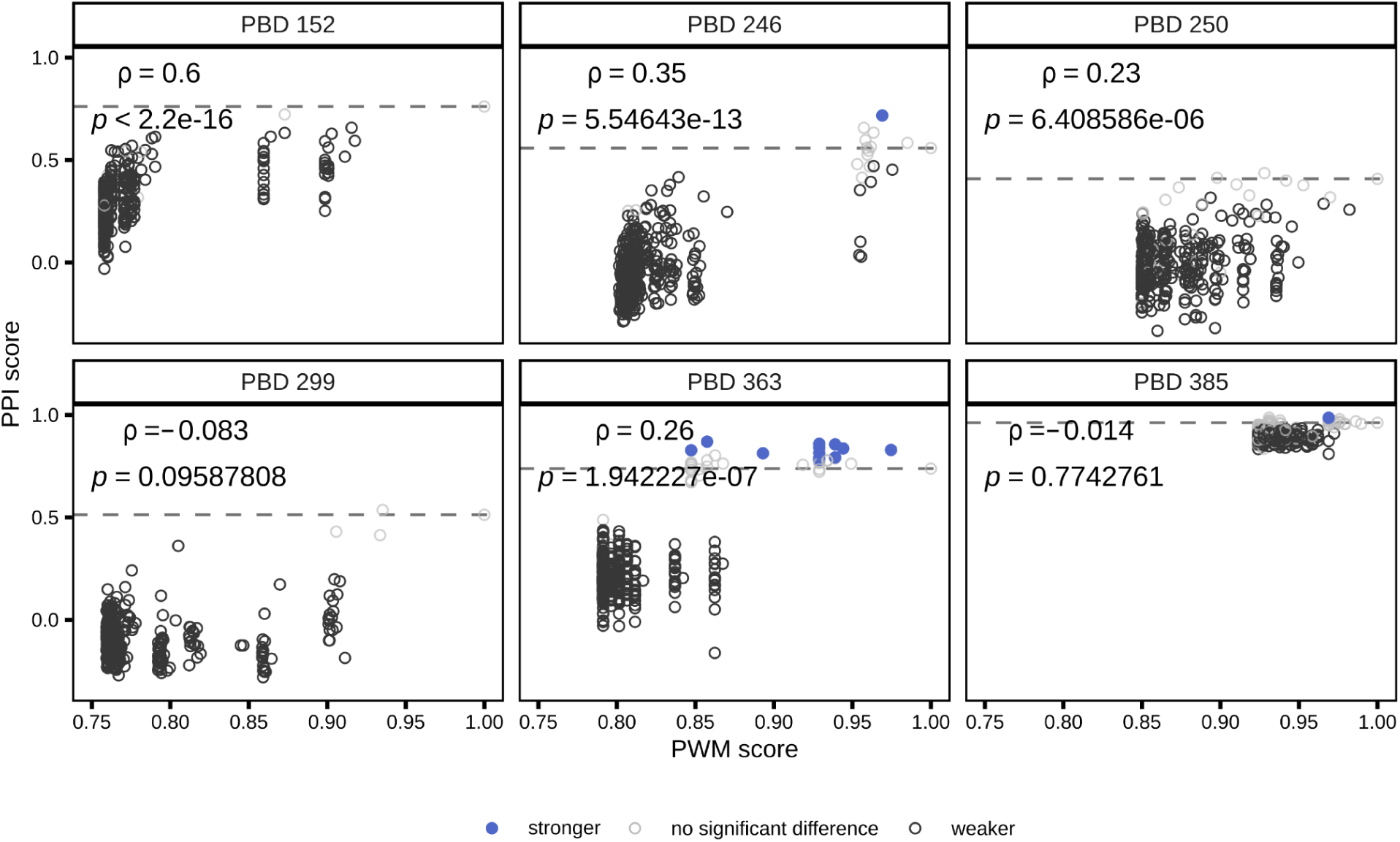
Comparison of PPI and PWM scores. Scatter plots showing the comparison between PPI score of double mutants (y-axis) and the score obtained by comparing the peptide sequences to the PWM extracted from the PRM-DB (x-axis), where a score of 1 is the complete agreement with the PWM (PWM_max_ peptide). The PBDs are labelled above each plot. Spearman’s correlation coefficients and their p-value are labelled for each comparison. The color of the data points indicates whether the peptide variants were found to bind significantly stronger or weaker than the PWM_max_ peptide to the labelled PBD. The dashed lines show the PPI score of the PWM_max_ peptides.

**Figure S6.**
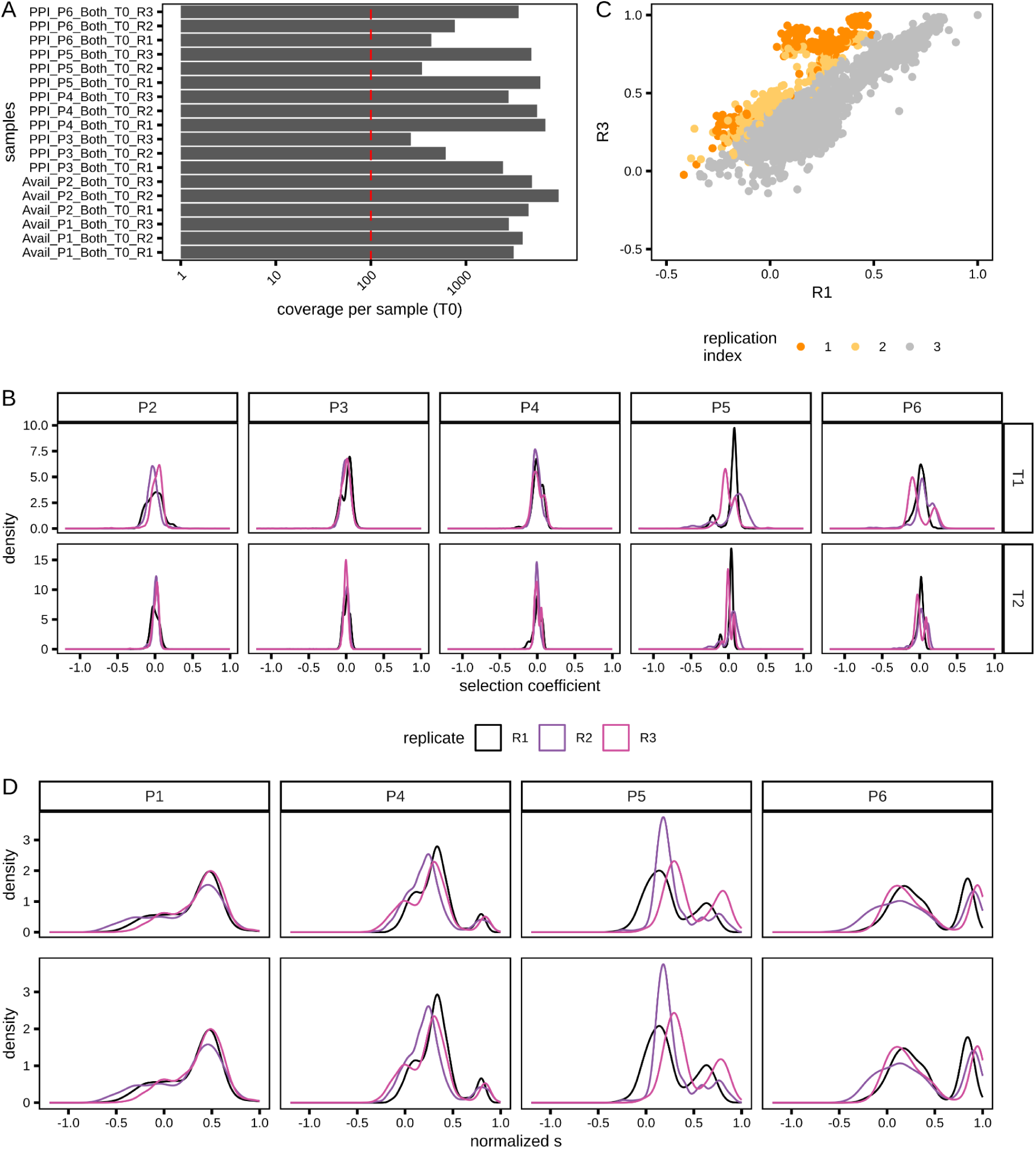
Processing of the read count of the second PCA bulk competition assay. **A.** Bar plot showing the coverage in reads of each sample at T0, where the red dashed line shows the 100X threshold. The y-axis described each sample with the assay type and the replicate. **B.** Density plots illustrating the selection coefficient distribution (see methods) computed between T0 and T1/T2 (rows) for the different pool described in Table S6 (columns) characterized for fitness effects (DMSO condition) **C.** Scatter plot comparing the normalised s of each DNA sequence between replicate 1 and replicate 3 of P5 in MTX condition. The color of the data points shows the replication index (see methods) of each DNA variant where the sequences with an index of 1 are filtered out of the dataset. **D.** Density plots illustrating the normalized s distribution (see methods) computed between T0 and T2 for the different pool (columns) characterized for PCA signal (MTX condition) before (top row) and after filtering (bottom row).

**Figure S7.**
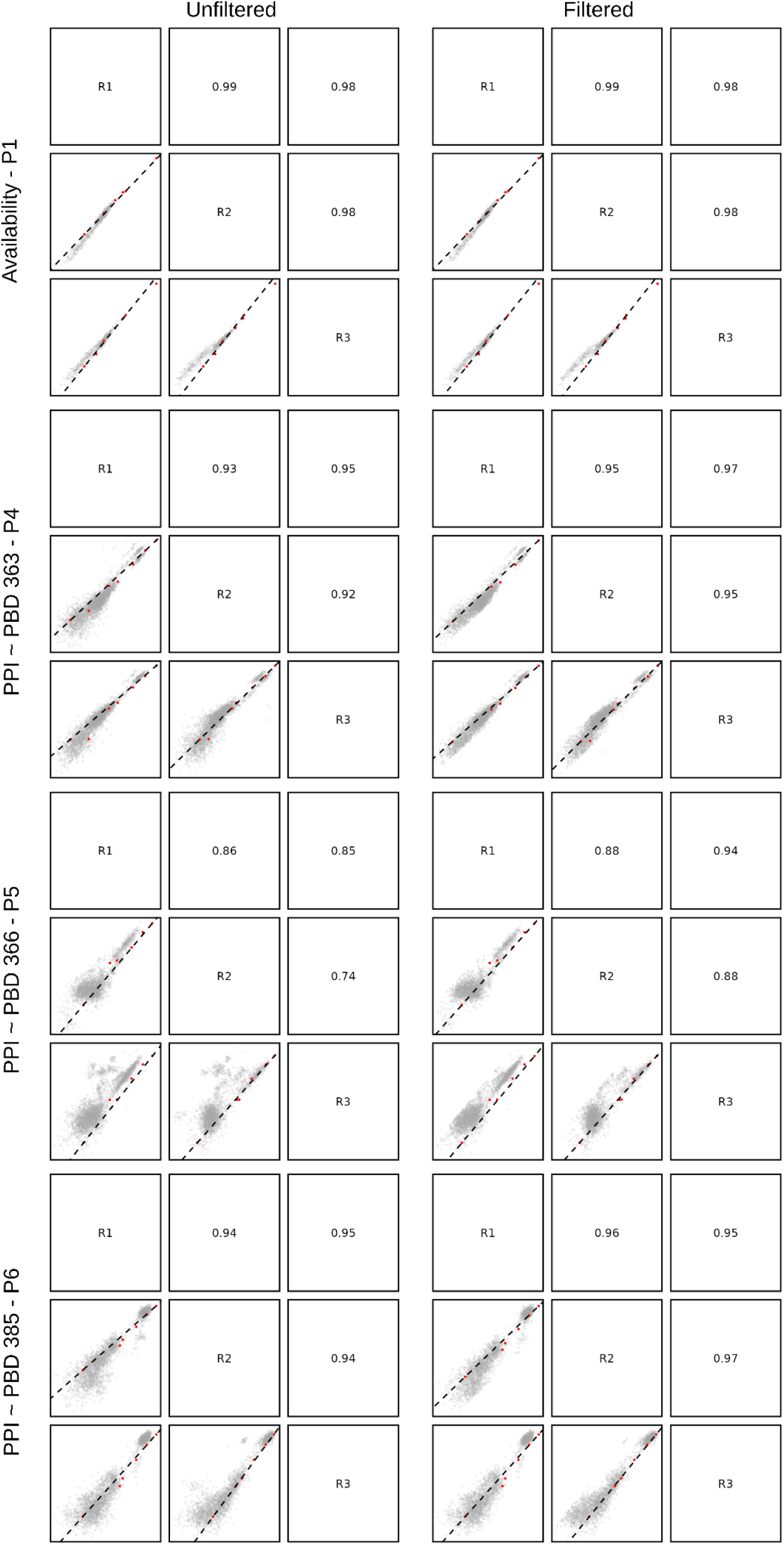
Evaluation of reproducibility and filtering of outliers for the second PCA bulk competition experiment. Scatter plots comparing the normalised s between replicates (R1, R2, R3) of the same pool (P1, P4, P5, P6) unfiltered (left) and after removing outliers (right) as explained in the methods. The replicates are identified on the diagonal of each grid and the Spearman’s correlation coefficients are shown on the mirror quadrant to each scatter plot (p-values < 2.2e-16). The red data points are the reference sequences spiked in the libraries. See Supplementary File 1 –Table S6 for the description of the composition of each pool.

**Figure S8.**
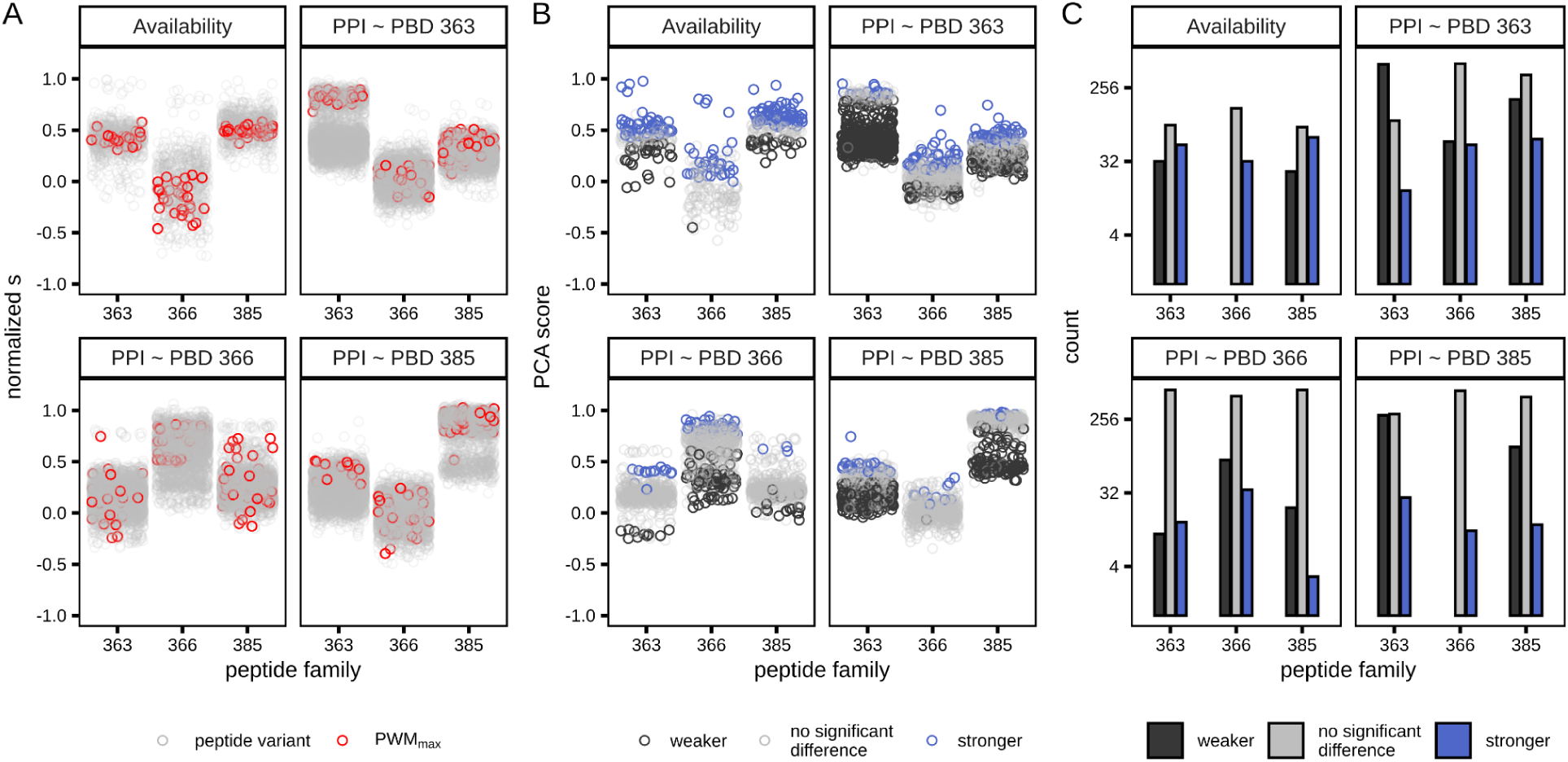
Identification of differences between PWM_max_ peptide and peptide variants in the second PCA bulk competition assay. For all panels, each plot represents a pool. The assay type and the PBD, for PPI assay, used in the pool are labelled on top of the plot. **A.** Distribution of the availability normalised s and the PPI normalised s for each DNA sequence associated with their peptide family on the x-axis. The colored data points show the synonymous sequence encoding the PWM_max_ peptide of each peptide family. **B.** The availability and PPI scores are shown for each peptide sequence associated with their peptide family on the x-axis. The colors of the data points represent the classification of each peptide as a statically stronger or weaker binder compared to the PWM_max_ peptide (Wilcoxon test, Benjamini-Hochberg correction, p-value < 0.05). **C**. Bar plots illustrating the number of weaker and stronger interacting peptide variants detected in each peptide family relative to the PWM_max_ peptide for availability and PPI assays (Wilcoxon test, Benjamini-Hochberg correction, p-value < 0.05).

**Figure S9.**
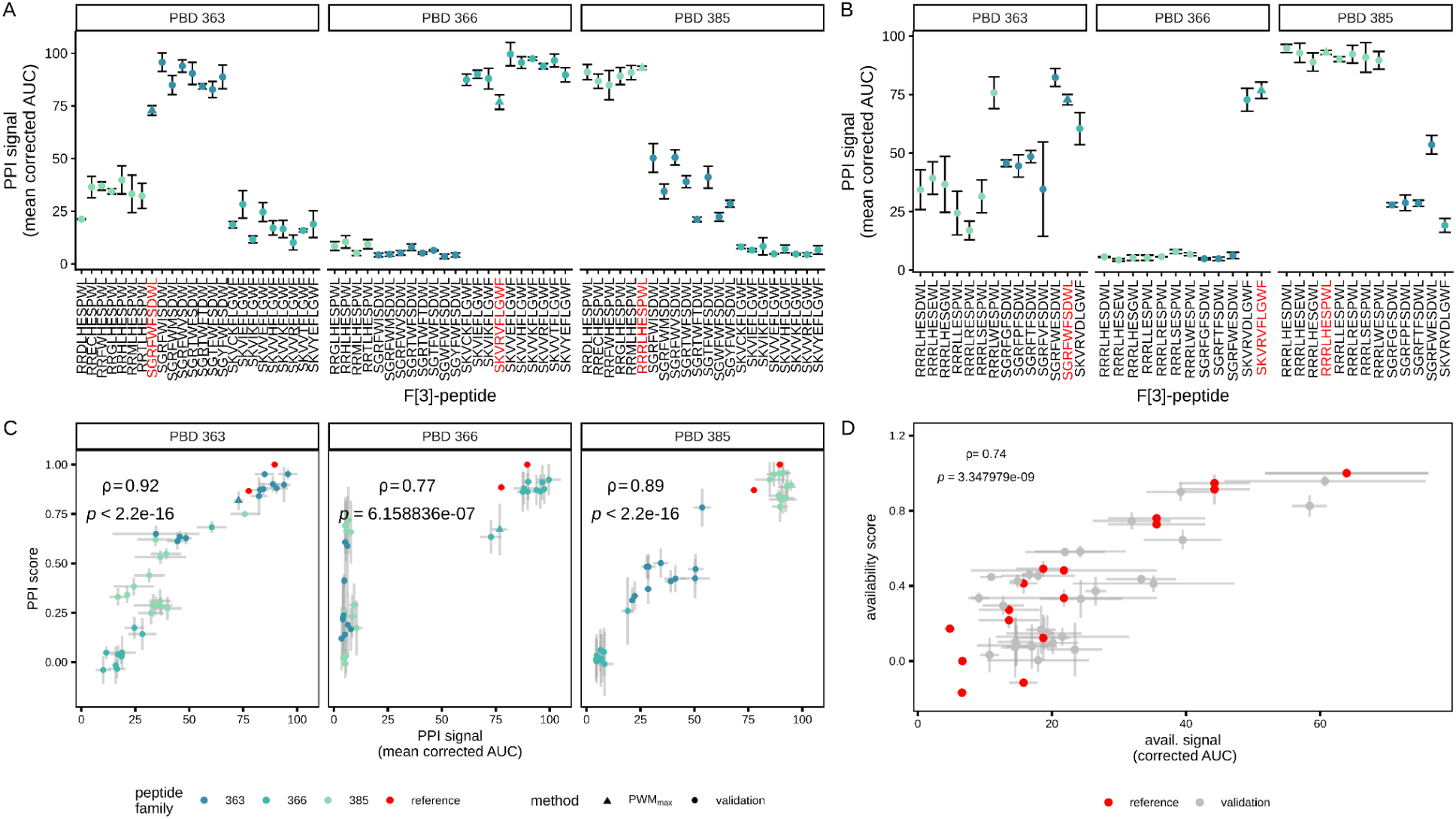
Small-scale validation of PPI and availability PCA bulk competition. **A.** The PBDs used for the PCA validation are labelled on top of the panels. For each peptide selected to validate an increased binding to a PBD (x-axis), the PPI signal obtained from growth curves measurements in triplicate is shown on the y-axis. The color of the data point represents the peptide families and the error bars show the standard deviation between replicates. The PWM_max_ peptides are represented as a triangle for comparison with the peptide variants and their sequences are colored in red. **B.** Same as panel A with peptides selected for validation of the change in specificity between peptide families and PBDs. **C.** Scatter plot comparing the PPI signal obtained from growth curve measurements for the validation peptides (x-axis) and the PPI score obtained from the bulk competition experiment for the PBDs identified on top of each plot. The Spearman’s correlation coefficient and p-value are labelled and error bars represent the standard deviation between replicates (n >= 3). The data point colors represent the peptide families and the red ones are reference peptides. The PWM_max_ peptides are represented as a triangle. **D.** Scatter plot comparing the availability signal obtained from growth curve measurements for the validation peptides (x-axis) and the availability score obtained from the bulk competition experiments. The Spearman’s correlation coefficient and p-values are labelled and error bars represent the standard deviation between replicates (n >= 3). The red data points are the reference sequences used in the bulk competition experiments.

**Figure S10.**
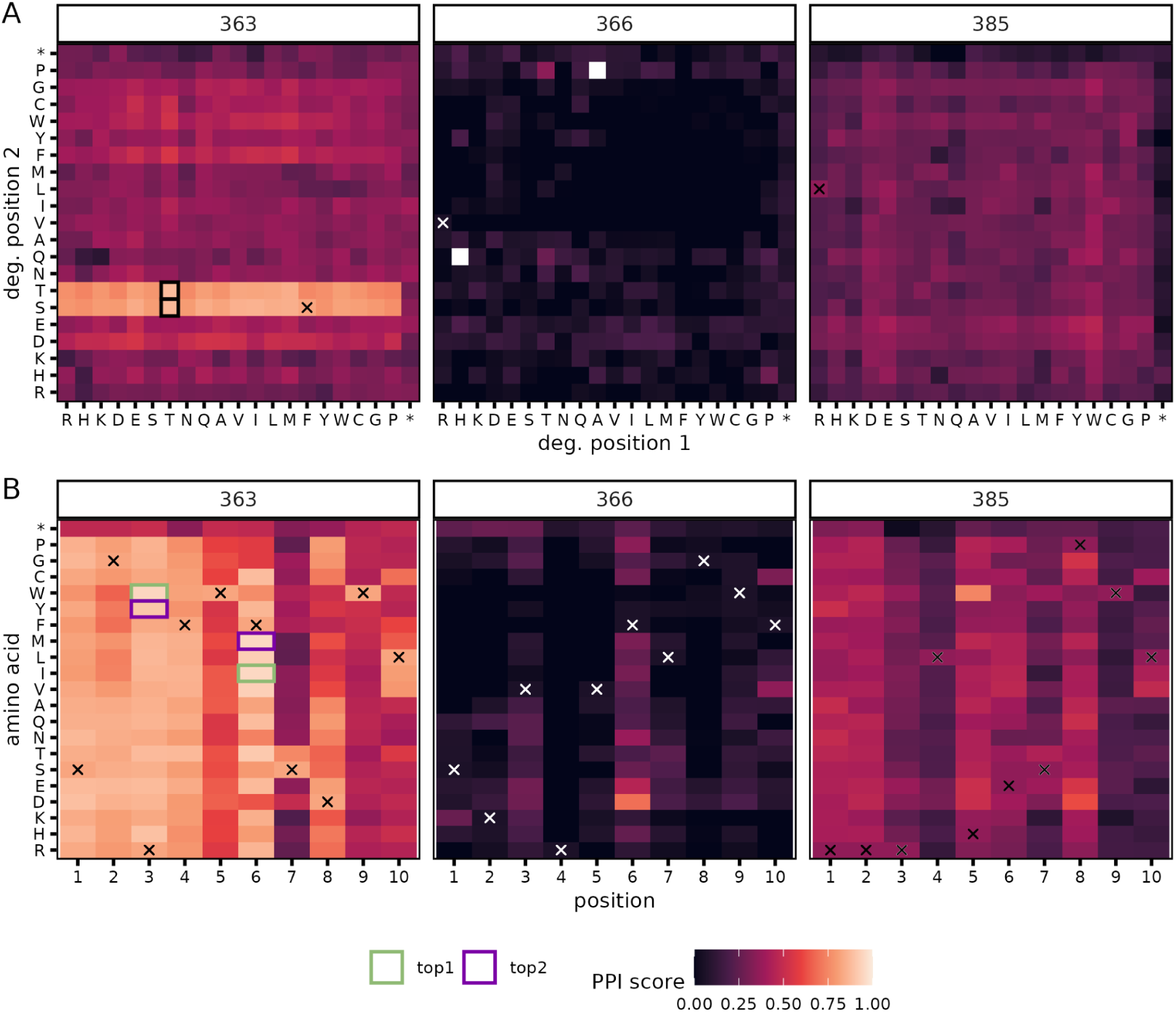
Specificity profile of PBD 363. The label on top of each plot is the peptide family tested for PPI with PBD 363. The heatmap color scale represents the PPI score and the crosses identify the PWM_max_ peptide sequence from the peptide family labelled above. Amino acid order follows their chemical properties (charged: R, H, K, D, E; polar: S, T, N, Q; hydrophobic: A, V, I, L, M, F, Y, W; special: C, G, P; stop: *). White tiles are missing data. **A.** Heatmap of the PPI score of each peptide variant present in the double mutant libraries. Each axis represents the amino acid encoded in one of the two degenerated positions in each peptide family. The black squares indicate the double mutant background used for peptide optimization of the 363 peptide family. **B.** Heatmap of the PPI score of each single mutant of the PWM_max_ peptide. The y-axis identifies the amino acid encoded in the positions of the peptide on the x-axis. The purple squares indicate the single mutants selected for peptide optimization of the 363 peptide family.

**Figure S11.**
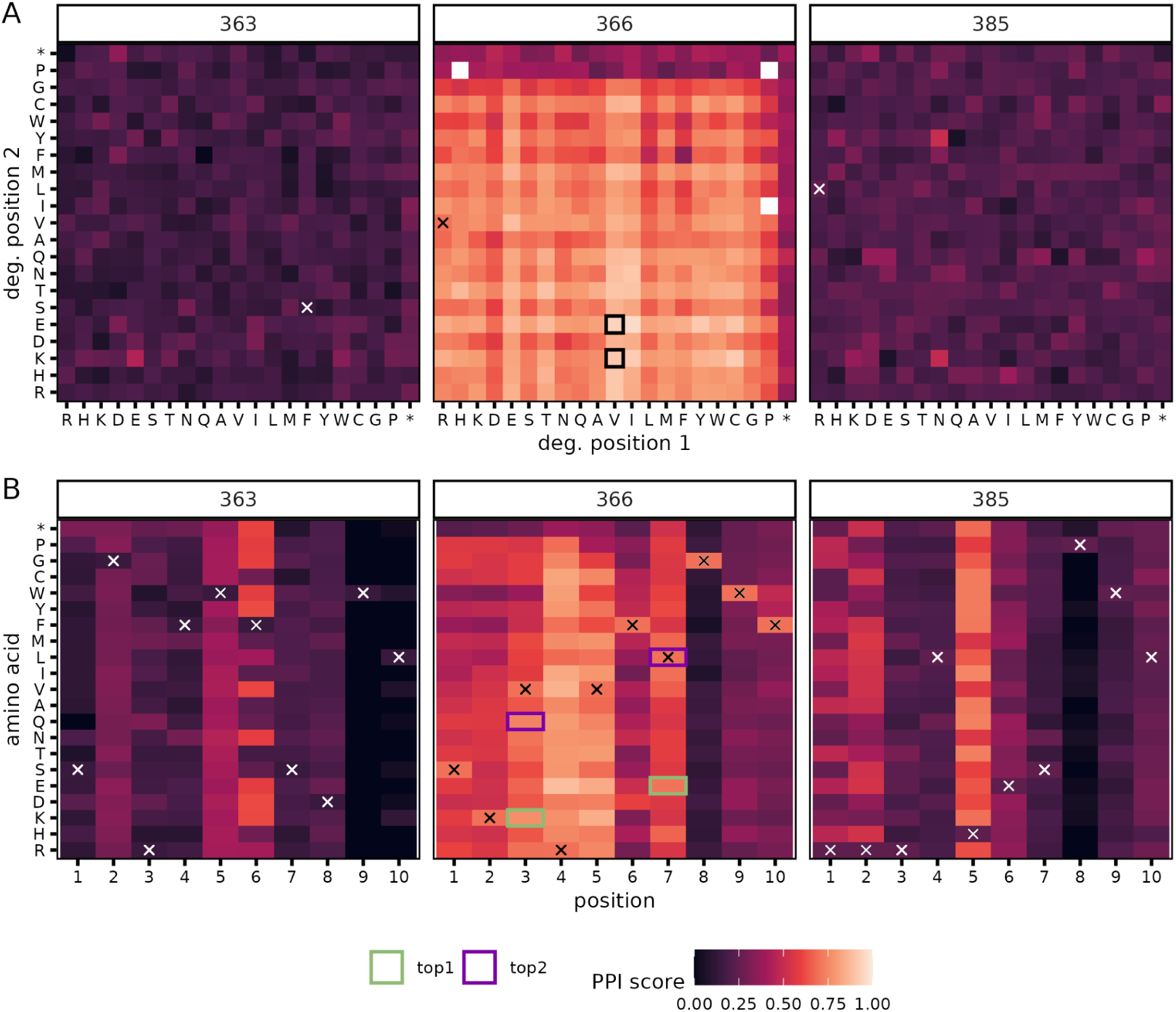
Specificity profile of PBD 366. The label on top of each plot is the peptide family tested for PPI with PBD 366. The color scale represents the PPI score and the crosses identify the PWM_max_ peptide sequence from the peptide family labelled above. Amino acid order follows their chemical properties (charged: R, H, K, D, E; polar: S, T, N, Q; hydrophobic: A, V, I, L, M, F, Y, W; special: C, G, P; stop: *). White tiles are missing data. **A.** Heatmap of the PPI score of each peptide variant present in the double mutant libraries. Each axis represents the amino acid encoded in one of the two degenerated positions in each peptide family. The black squares indicate the double mutant background used for peptide optimization of the 366 peptide family. **B.** Heatmap of the PPI score of each single mutant of the PWM_max_ peptide. The y-axis identifies the amino acid encoded in the positions of the peptide on the x-axis. The purple squares indicate the single mutants selected for peptide optimization of the 366 peptide family.

**Figure S12.**
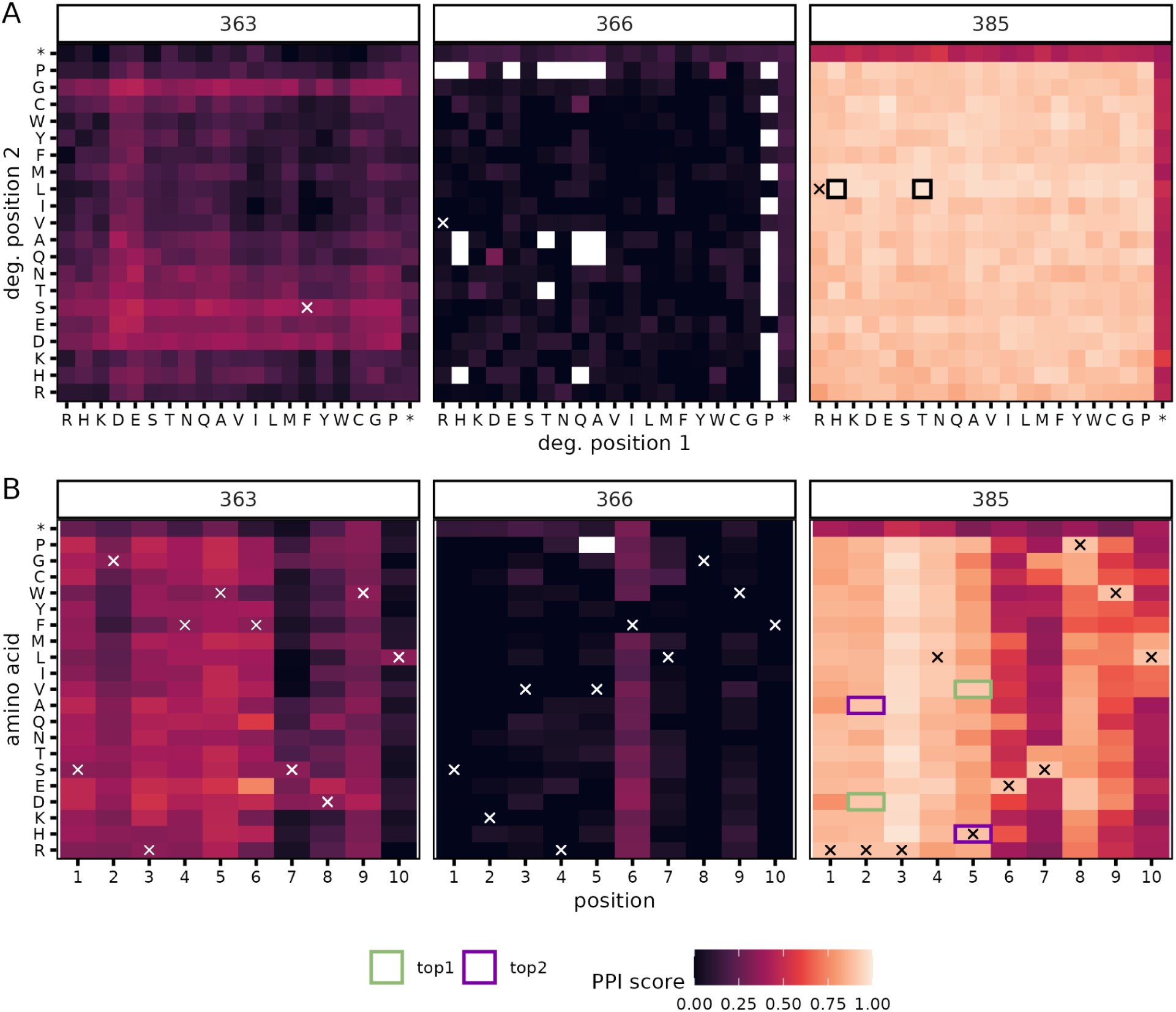
Specificity profile of PBD 385. The label on top of each plot is the peptide family tested for PPI with PBD 385. The color scale represents the PPI score and the crosses identify the PWM_max_ peptide sequence from the peptide family labelled above. Amino acid order follows their chemical properties (charged: R, H, K, D, E; polar: S, T, N, Q; hydrophobic: A, V, I, L, M, F, Y, W; special: C, G, P; stop: *). White tiles are missing data. **A.** Heatmap of the PPI score of each peptide variant present in the double mutant libraries. Each axis represents the amino acid encoded in one of the two degenerated positions in each peptide family. The black squares indicate the double mutant background used for peptide optimization of the 385 peptide family. **B.** Heatmap of the PPI score of each single mutant of the PWM_max_ peptide. The y-axis identifies the amino acid encoded in the positions of the peptide on the x-axis. The purple squares indicate the single mutants selected for peptide optimization of the 385 peptide family.

**Figure S13.**
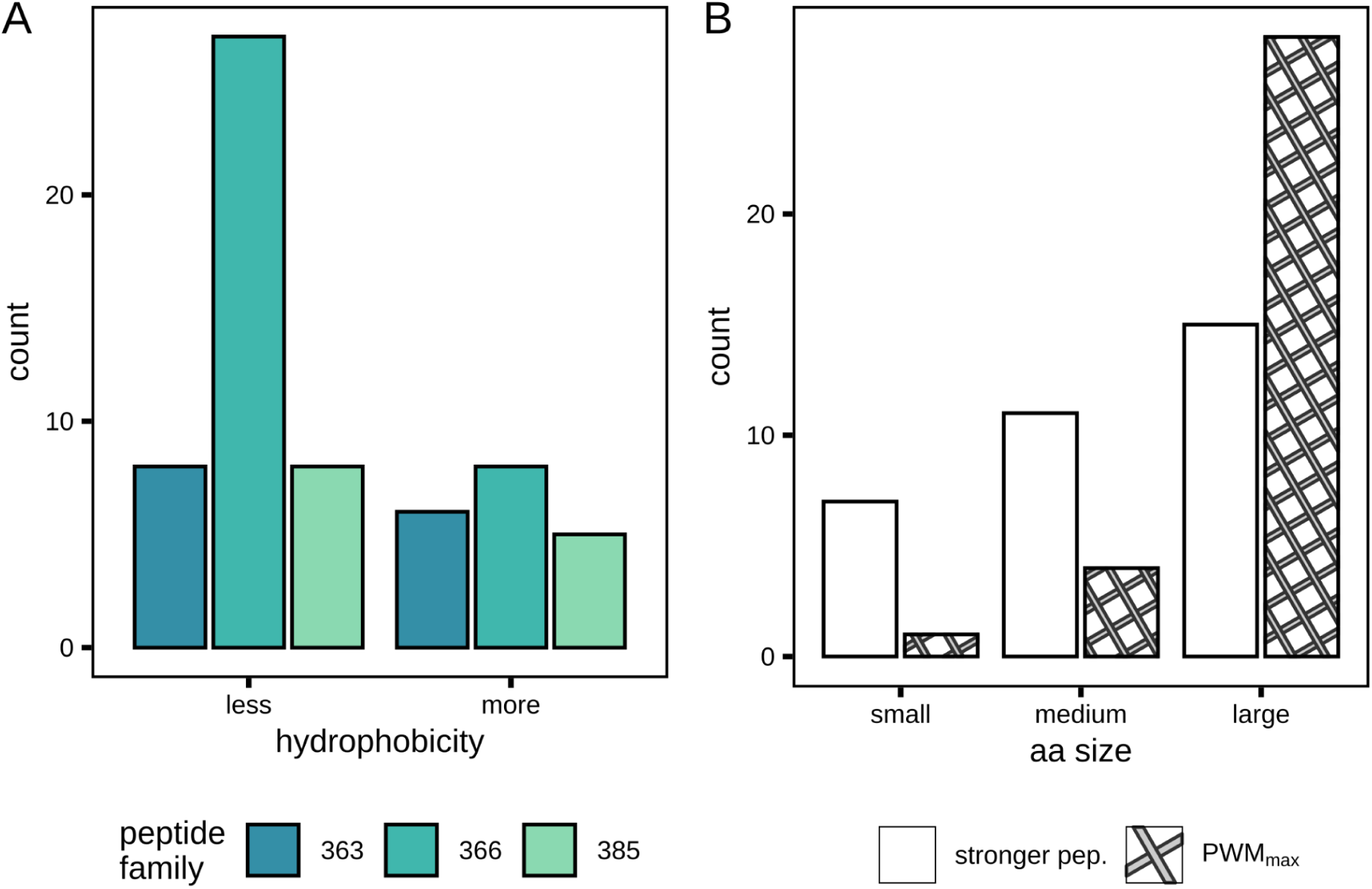
Amino acid properties of stronger interacting peptide variants. **A.** Bar plot representing the number of stronger interacting peptide variants which are less or more hydrophobic (scored using the Cornell scale) than the PWM_max_ peptide for each peptide family. **B.** Bar plot showing the distribution of size of the reference amino acid encoded in the PWM_max_ peptides and of the amino acids leading to a gain of interaction. Amino acid sizes are attributed as small: G, A, S; medium: V, P, C, T, D, N; large: F, Y, H, L, M, Q, W, I, R, E, K.

**Figure S14.**
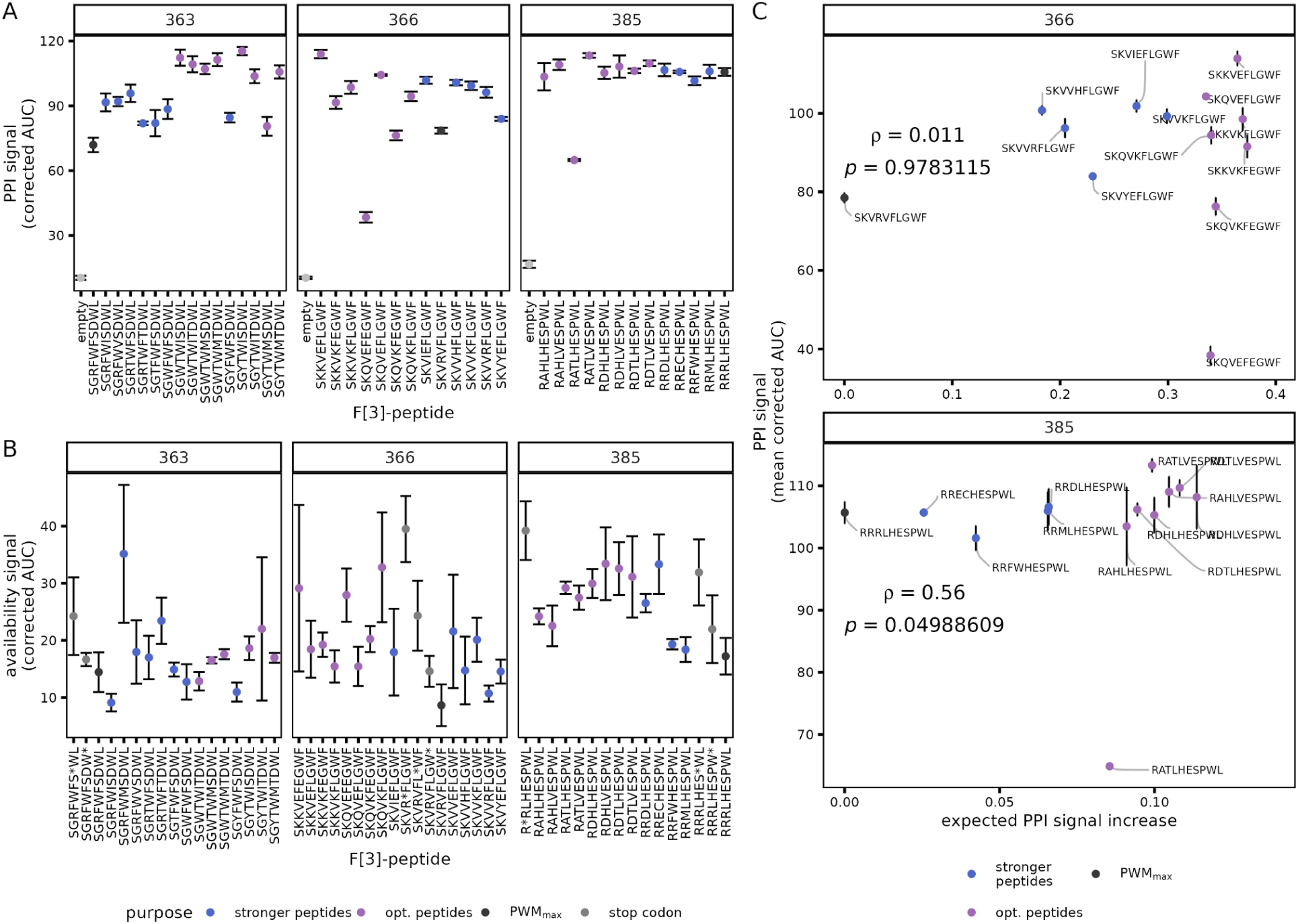
PPI and availability characterization of optimized peptides. Each data point represents the mean signal for peptides tested in PCA small-scale assays (n = 3) for **A**. the PPI assay with the PBD labelled above the plot and **B.** the availability assay for the peptides in the peptide family labelled above the plot. **C.** Scatter plot showing the comparison between the PPI signal and the expected PPI signal increase of the stronger and optimized peptides relative to the PWM_max_ peptide based on the PPI scores obtained in the bulk competition for the 366 (top) and 385 (bottom) peptide families. The peptide sequences are labelled for each data point. Spearman’s correlation coefficient and p-value are labelled. For all panels, error bars show the standard deviations between triplicates and the data point color identifies the peptide type.

**Figure S15.**
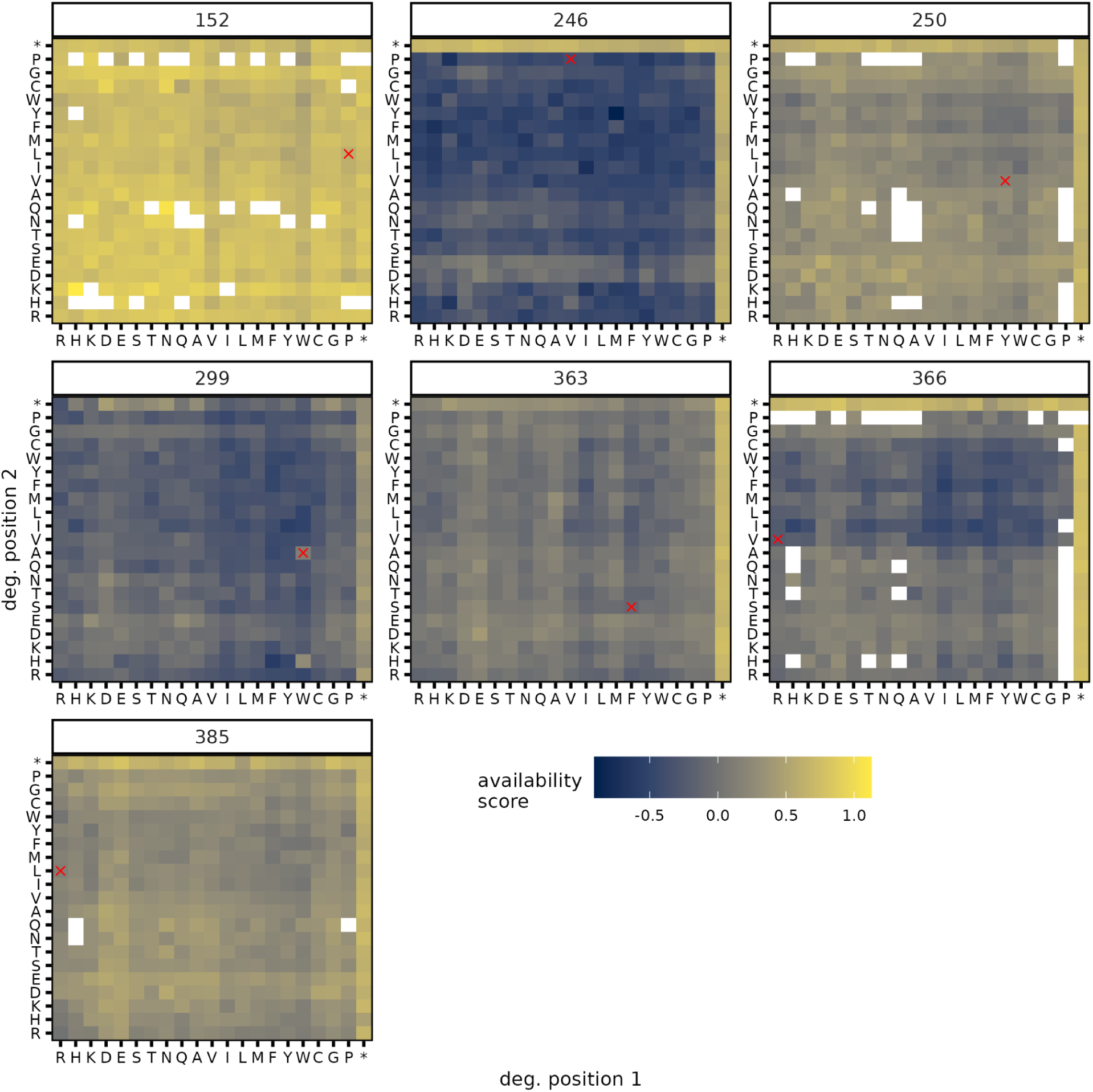
Binding availability detected by DHFR PCA bulk competition of double mutants. The peptide families are labelled on the top of each panel. The color scale of the heatmaps represents the binding availability score and the pairs of mutated residues are labelled on the axis. Amino acid order follows their chemical properties (charged: R, H, K, D, E; polar: S, T, N, Q; hydrophobic: A, V, I, L, M, F, Y, W; special: C, G, P; stop: *). The cross highlights the maximal binding peptide defined by the PWMs. White tiles are missing data.

**Figure S16.**
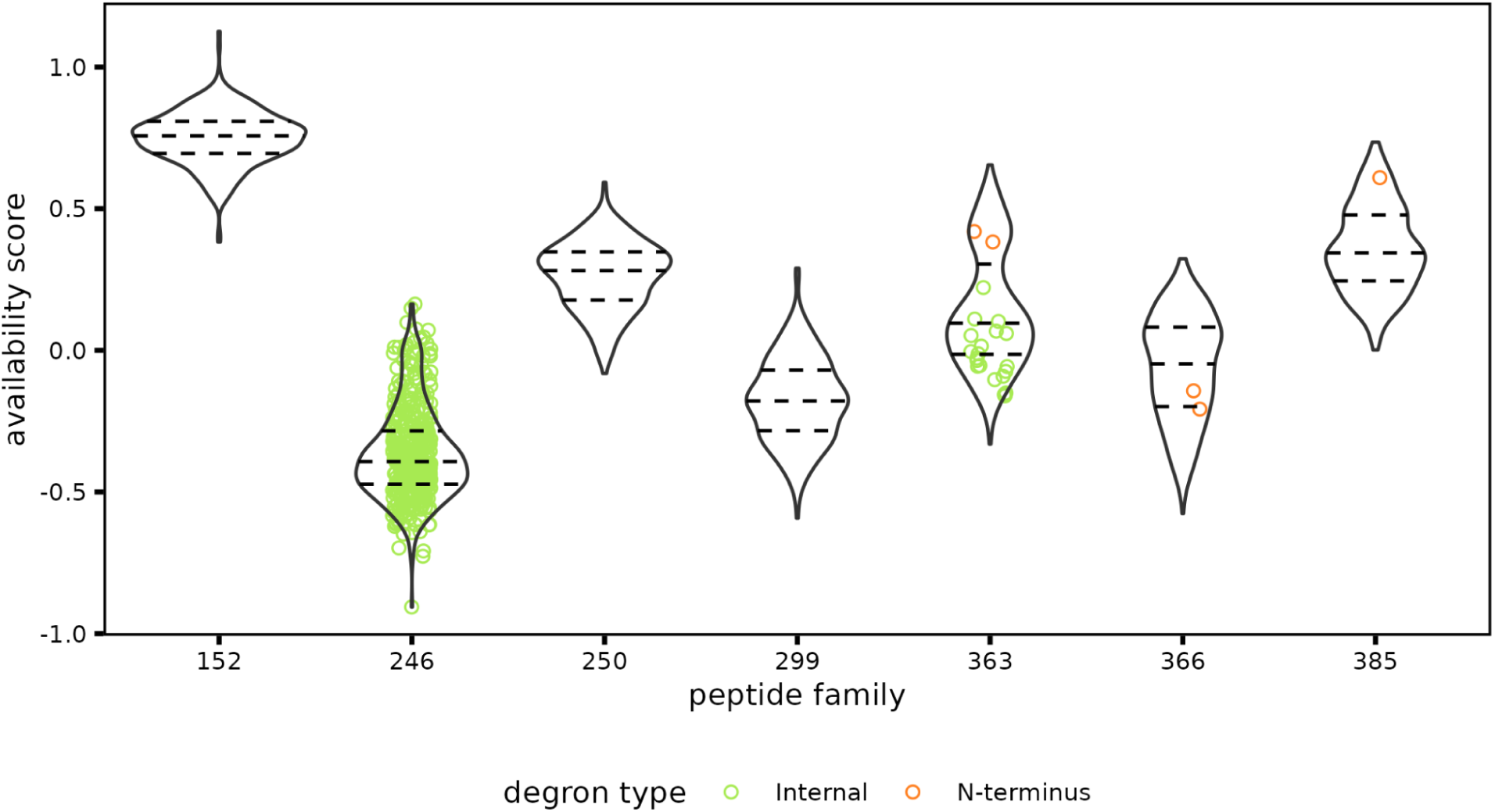
Degron presence in peptide variants compared to binding availability. Violin plot showing the distribution of availability signals of single and double mutants of PWM_max_ peptides. The dashed lines show the 25th, 50th and 75th quantiles of each distribution. The colored data points highlight the peptide variants in which a known yeast degron was found (Szulc et al., 2024).

**Figure S17.**
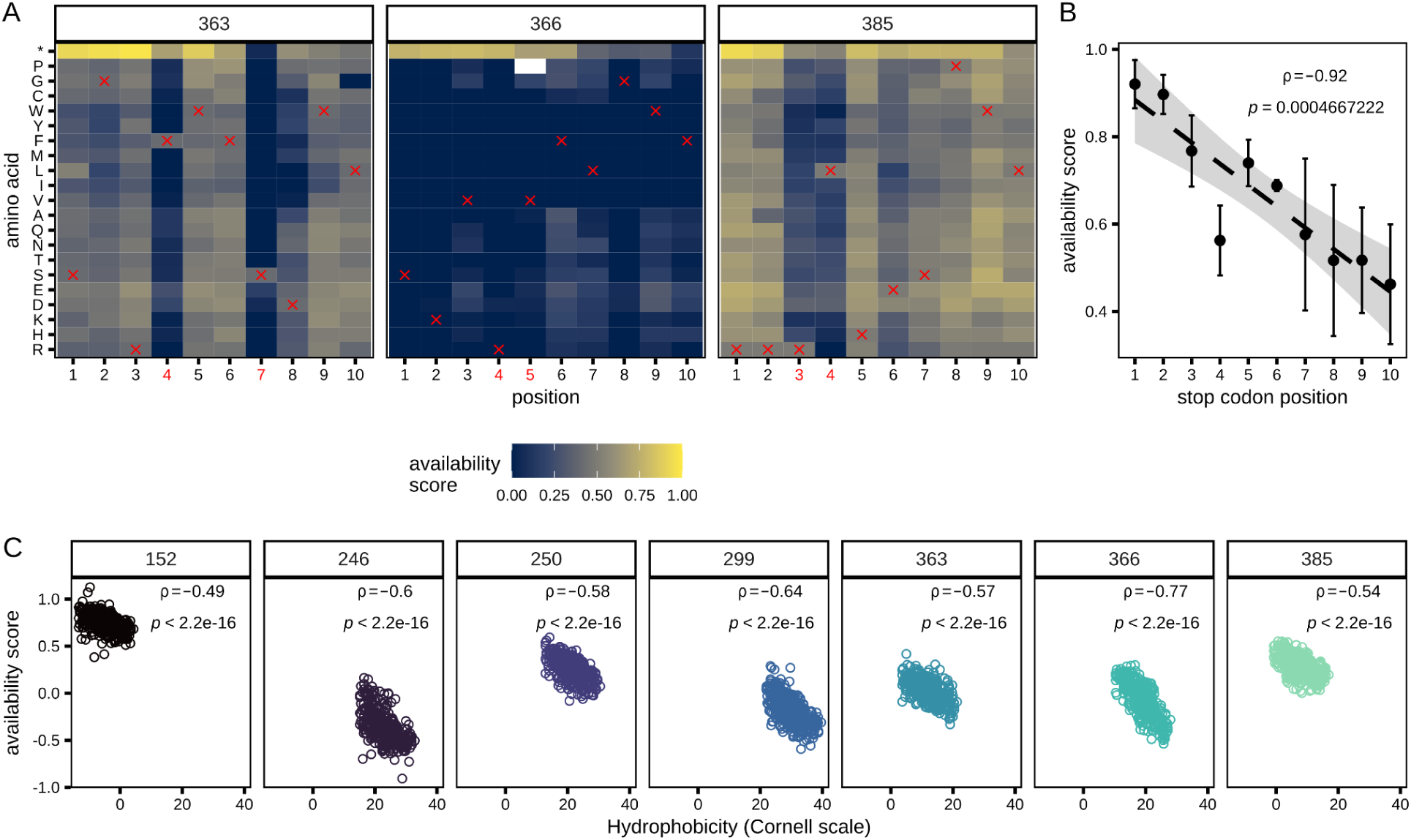
Availability and hydrophobicity of peptide variants. **A.** The label on top of each heatmap is the peptide family. The color scale represents the availability score and the crosses identify the PWM_max_ peptide sequence from the peptide family labelled above. Amino acid order follows their chemical properties (charged: R, H, K, D, E; polar: S, T, N, Q; hydrophobic: A, V, I, L, M, F, Y, W; special: C, G, P; stop: *). White tiles are missing data. The x-axis identifies the amino acid position in the peptide. The positions identified in red are the degenerated positions in the double mutant libraries and the data shown for those positions are extracted from the first PCA experiment while the black labelled positions data were obtained in the second PCA competition experiment. **B.** Relation between the stop codon position in PWM_max_ peptides and their availability signal. Data point represents the mean of availability score and the error bars show the maximal and minimal value of availability score per position. Spearman’s correlation coefficient and p-value between the mean availability value and the position are labelled. **C.** Isolated comparison of availability score and hydrophobicity per peptide family. The label on top of each scatter plot is the peptide family. Spearman’s correlation coefficients and p-values are labelled.

**Figure S18.**
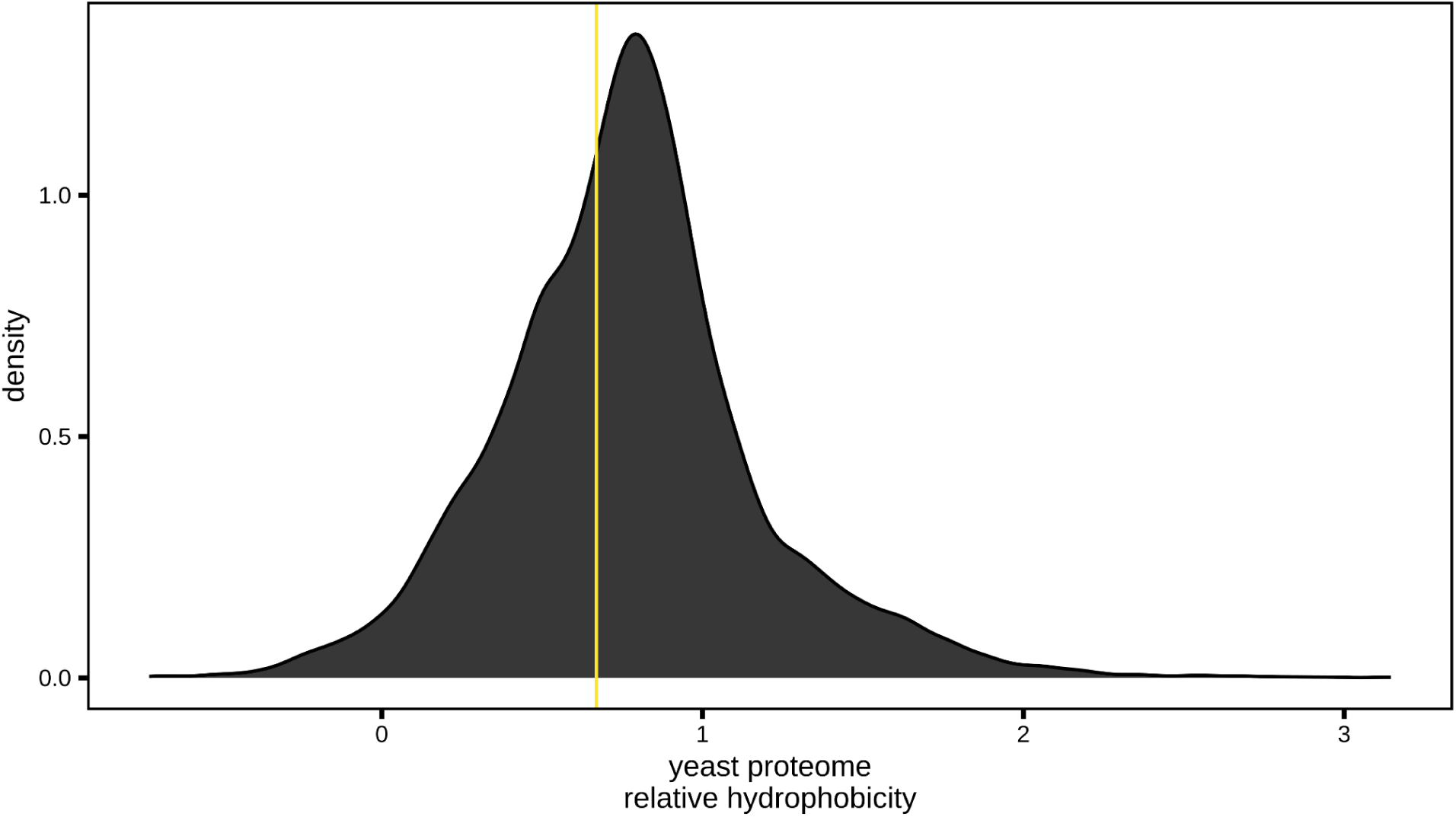
Relative hydrophobicity of the yeast proteome. Density plot showing the relative Cornell hydrophobicity (Karplus, 1997), mean hydrophobicity per position, of all yeast proteins. The yellow line highlights the relative Cornell hydrophobicity of the YFP-F[1,2].

